# A widespread toxin-antitoxin system exploiting growth control via alarmone signalling

**DOI:** 10.1101/575399

**Authors:** Steffi Jimmy, Chayan Kumar Saha, Constantine Stavropoulos, Sofia Raquel Alves Oliveira, Tatsuaki Kurata, Alan Koh, Albinas Cepauskas, Hiraku Takada, Tanel Tenson, Henrik Strahl, Abel Garcia-Pino, Vasili Hauryliuk, Gemma C. Atkinson

## Abstract

Under stressful conditions, bacterial RelA-SpoT Homologue (RSH) enzymes synthesise the alarmone (p)ppGpp, a nucleotide messenger. (p)ppGpp rewires bacterial transcription and metabolism to cope with stress, and at high concentrations inhibits the process of protein synthesis and bacterial growth to save and redirect resources until conditions improve. Single domain Small Alarmone Synthetases (SASs) are RSH family members that contain the (p)ppGpp synthesis (SYNTH) domain, but lack the hydrolysis (HD) domain and regulatory C-terminal domains of the long RSHs such as Rel, RelA and SpoT. We have discovered that multiple SAS subfamilies can be encoded in broadly distributed conserved bicistronic operon architectures in bacteria and bacteriophages that are reminiscent of those typically seen in toxin-antitoxin (TA) operons. We have validated five of these SASs as being toxic (toxSASs), with neutralisation by the protein products of six neighbouring antitoxin genes. The toxicity of *Cellulomonas marina* ToxSAS FaRel is mediated by alarmone accumulation combined with depletion of cellular ATP and GTP pools, and this is counteracted by its HD domain-containing antitoxin. Thus, the ToxSAS-antiToxSAS system is a novel TA paradigm comprising multiple different antitoxins that exemplifies how ancient nucleotide-based signalling mechanisms can be repurposed as TA modules during evolution, potentially multiple times independently.

## INTRODUCTION

Bacteria encounter a variety of different environmental conditions during their life cycles, to which they need to respond and adapt in order to survive. This can include slowing down their growth and redirecting their metabolic resources during nutritional stress, until conditions improve and the growth rate can increase. One of the main signals that bacteria use for signalling stress are the alarmone nucleotides ppGpp and pppGpp, collectively referred to as (p)ppGpp^1^. At high concentrations (p)ppGpp is a potent inhibitor of bacterial growth^2^, targeting transcription, translation and ribosome assembly^1^. (p)ppGpp is produced and degraded by proteins of the RelA/SpoT homologue (RSH) superfamily, named after the two *Escherichia coli* representatives – multi-domain ‘long’ RSH factors RelA and SpoT^3^. In addition to long RSHs, bacteria can encode single-domain RSHs: Small Alarmone Synthetases (SAS) and Small Alarmone Hydrolases (SAH).

It is currently unknown why some bacteria carry multiple SASs and SAHs, which can belong to many different subfamilies. Conservation of gene order through evolution can reveal potentially interacting proteins and shed light on the cellular role of proteins^4^. Therefore, we developed a computational tool – FlaGs, standing for Flanking Genes^5^ – for analysing the conservation of genomic neighbourhoods, and applied it to our updated database of RSH sequences classified into subfamilies. Surprisingly, we find that some subfamilies of SAS can be encoded in apparently bi- (and sometimes tri-) cistronic, often overlapping, conserved gene architectures that are reminiscent of toxin-antitoxin (TA) loci^6^. The potential for SAS toxicity is supported by the observation that when (p)ppGpp is over-produced – for example, if synthesis by RelA is not balanced by hydrolysis by SpoT – the alarmone becomes toxic and inhibits growth^7^.

The first direct evidence that RSH toxicity *per se* might be a *bona fide* function of some SASs was provided by Dedrick and colleagues^8^. They showed that gp29, a SAS encoded by the mycobacterial Cluster N bacteriophage Phrann is exceedingly toxic to *M. smegmatis.* This toxicity is countered by co-expression of its neighbouring gene (gp30) – a proposed inhibitor of the SAS. Neither the molecular mechanism of gp29-mediated toxicity nor its neutralisation by gp30 are known. The gp29-mediated abrogation of growth is proposed to be a defence mechanism against co-infection by other bacteriophages, such as Tweety and Gaia^8^.

The regulatory interplay between gp29 and gp30 is typical of that seen in toxin-antitoxin (TA) systems. When expressed, the protein toxin abolishes bacterial growth – and its toxicity can be efficiently countered by the protein or RNA antitoxin. Known toxins can act in a number of ways^6^, commonly by targeting translation by cutting or modifying the ribosome, translation factors, tRNAs or mRNAs. Similarly, antitoxins counteract the toxins through different mechanisms^6^: through base-pairing of the antitoxin RNA with the toxin mRNA (type I TA systems), direct protein-protein inhibition (type II), inhibition of the toxin by the antitoxin RNA (type III), or by indirect nullification of the toxicity (type IV).

In this study we have uncovered the evolutionary diversity of SAS-based toxin (toxSAS) TA systems using sensitive *in silico* sequence searching and gene neighbourhood analysis. We have experimentally validated five SAS subfamilies as belonging to *bona fide* TA systems and demonstrated through mutagenesis that the toxicity of SASs is strictly dependent on a functional (p)ppGpp synthetase active site. Of our six identified antitoxins, five are strictly specific in counteracting only their cognate toxSAS and one other can universally neutralize all of the toxSASs. This antitoxin encodes a (p)ppGpp degrading enzyme – SAH – and acts as a type IV antitoxin degrading the molecular product of toxSAS synthetic activity.

## MATERIAL AND METHODS

### Identification and classification of RSH sequences across the tree of life

Predicted proteomes from 24072 genomes were downloaded from NCBI, selecting one representative genome per species for Archaea, Bacteria, Eukaryotes and all Viruses. The sequences from our previous RSH database^3^ were extracted on a subfamily basis, aligned with MAFFT v7.313^9^ and hidden Markov models (HMMs) were made with HMMer v3.1b2^10^. All genomes were scanned with the HMMs to identify RSH family members and classify them into subfamilies with E value cut-off thresholds that were previously determined^3^: E^-4^ and E^-5^ for SYNTH and HD domains, respectively. HMMs of the HD and SYNTH domains were used to determine the (p)ppGpp synthesising and hydrolysing domains present in the 35615 identified sequences. Sequences, taxonomy of the source organism, domain composition and subfamily memberships were stored in a MySQL database. To update the classifications with novel subfamilies, phylogenetic analysis was carried out of RSHs from representative genomes based on taxonomy (one representative per genus). Three data sets of sequences (long RSHs, SASs and SAHs) were extracted and aligned as above. Phylogenetic analysis was carried out with FastTree v2.1.10^11^ after removing positions containing more than 50% gaps, and any extremely divergent proteins that could not be confidently aligned. The three resulting trees and alignments were examined by eye with FigTree v1.4.2 and Aliview v1.2.0^12^ to identify groups that are appear to be distinct, that is, are comprised of mostly orthologues, have a distinct domain structure, and, ideally, have strong support for monophyly. Eight of our subfamilies are paraphyletic in that they contain one or monophyletic groups nested within their diversity: MixSpo, AaRel, CapRel, FpRel, FunRel, MixRel, PRel3, and Rel. To further resolve relationships among subfamilies, trees were then made with RaxML v8.2.10^13^ and IQ-TREE v1.6.6^14^ on the Cipres Science Gateway v 3.3 portal^15^, excluding divergent sequences that could not be assigned to subfamilies in the FastTree tree. For Maximum Likelihood phylogenetic analyses, we used the LG model of substitution, which is the best-fit model for our dataset, as predicted by IQ-TREE. RAxML was run with 100 bootstrap replicates to give a value (maximum likelihood boostrap, MLB percentage) for how much of the input alignment supports a particular branch in the tree topology. In the case of IQ-TREE, the ultrafast bootstrapping (UFB) approximation method was used to compute support values for branches out of 1000 replicates. Trees from RAxML and IQ-TREE were visualized with FigTree and subfamilies were inspected for whether they contain mostly orthologues with at least moderate (>60%) bootstrap support. Overall, we could classify sequences into 13 subfamilies of long RSHs, 30 subfamilies of SASs and 11 subfamilies of SAHs. The sequences of each subfamily were aligned and used to make HMMs, as above. The 35615 sequences in the MySQL database were re-scanned with the updated subfamily HMMs and the database was updated, as reproduced here as two Excel files of all sequences (**Supplementary Table S1**), and subfamily distribution across taxonomy (**Supplementary Table S2**).

### Phylogenetic analysis of RSH subfamily representatives

To make the representative trees of **Figure 1**, we selected taxa from the RSH database to sample broadly across the tree of life, and cover all subfamilies of RSHs. We used a Python script to select 15 representatives per SAS or SAH subfamily, 145 representatives for the almost universal bacterial protein Rel, and 80 representatives for RelA and SpoT, based on taxonomy of the RSH-encoding organism. The script calculates the total number of unique names on each taxonomic level (e.g., phylum, class, order, genus and species) and optimises selection of representative sequences accordingly, in order to sample taxonomy as broadly as possible within that subfamily. The (p)ppGpp hydrolase (HD) domain-containing dataset and the (p)ppGpp synthetase (SYNTH) domain-containing dataset were separately aligned with MAFFT with the L-ins-i strategy, and alignment positions with >50% gaps were removed. After alignment curation, our HD domain-containing alignment contained 698 amino acid positions from 519 sequences, and the SYNTH domain-containing alignment contained 699 amino acid positions from 722 sequences (**Supplementary Text S1**). The two domain alignments were used for Maximum Likelihood phylogenetic analysis using RaxML and IQ-TREE as described above.

**Figure 1.**
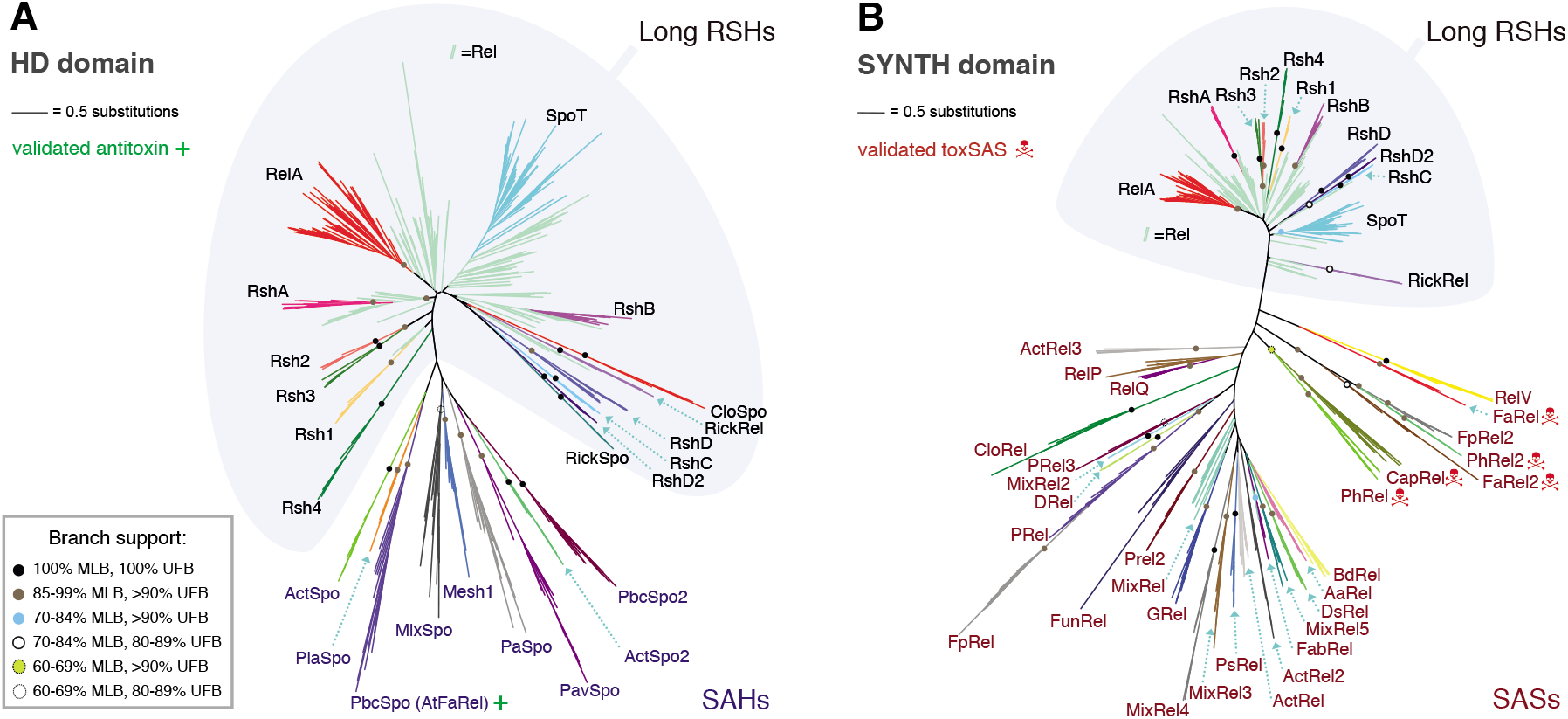
Maximum likelihood phylogenies of the (p)ppGpp hydrolase (A) and synthetase (B) domains. Trees were generated from RaxML and IQ-TREE analyses of alignments of representatives across the RSH family with (**A**) the (p)ppGpp hydrolase (HD) domain-containing dataset (698 amino acid positions, 519 sequences), and (**B**) the ppGpp synthetase (SYNTH) domain-containing dataset (699 amino acid positions, 722 sequences). Shading behind the branches indicates the boundary between multi-domain type (long) RSHs and single domain (small) RSHs. The long RSH groups also contain members that seem to have secondarily lost domains through evolution to become single domain (members of the RickSpo and RickRel groups). The inset box shows the legend for subfamily and intersubfamily support, support values within subfamilies, and those that are less that 60% MLB are not shown. Branch length is proportional to the number of substitutions per site (see scale bar). The red skull and crossbones symbol indicates those subfamilies of SASs that we have confirmed with toxicity neutralisation assays to contain toxSASs. The SAH group PbcSpo that we have found contains an antitoxin is indicated with a green plus sign. Alignments used for phylogenetic analysis, and trees with all branch support values are available in **Supplementary Text 1**.

### Analysis of gene neighbourhood conservation

Our Python tool FlaGs^5^ was used find conserved genomic architectures, using the NCBI accession numbers of representative RSH subfamily members (one per genus) as input. The legend files for the gene cluster numbers in all conservation figures in this paper are found in **Supplementary Table S3**.

### Prediction of prophage-encoded RSHs

To detect if bacterial SAS or SAH genes are located in bacteriophage-like sequence regions, we used the PHASTER URLAPI^16^. To create the input nucleotide data sets, we made a pipeline that takes the nucleotide sequence containing the four up and downstream genes around each SAS or SAH genes that are present in our RSH Database. The resulting PHASTER predictions are found in **Supplementary Table S4**.

### Toxicity neutralisation assays

Toxicity-neutralisation assays were performed on LB medium (Lennox) plates (VWR). *E. coli* BW25113 strains transformed with pBAD33 (encoding toxins) and pKK223-3 (encoding antitoxins) were grown in liquid LB medium (BD) supplemented with 100 μg/ml carbenicillin (AppliChem) and 20 μg/ml chloramphenicol (AppliChem) as well as 1% glucose (repression conditions). Serial ten-fold dilutions were spotted (5 μl per spot) on solid LB plates containing carbenicillin and chloramphenicol in addition to either 1% glucose (repressive conditions), or 0.2% arabinose combined with 1 mM IPTG (induction conditions). Plates were scored after an overnight incubation at 37 °C.

### Growth assays

Growth assays were performed in liquid MOPS minimal medium (1x MOPS mixture (AppliChem), 0.132 M K_2_HPO_4_ (VWR Lifesciences), 1mg/ml thiamine (Sigma), 0.1% casamino acids (VWR Lifesciences) and the carbon source – either 0.5% glycerol (VWR Chemicals) or 1% glucose). The media was supplemented with carbenicillin and chloramphenicol. Overnight cultures were grown in MOPS medium supplemented with 1% glucose at 37 °C. The cultures were diluted to a final OD_600_ of 0.01 in MOPS medium supplemented with 0.5% glycerol, 0.2% arabinose and 1mM IPTG. Growth was then monitored using a Bioscreen C Analyzer (Oy Growth Curves Ab Ltd) at 37 °C for 10 hours.

The experimental procedures for construction of plasmids, microscopy, quantification of nucleotide pools by TLC and HPLC, protein expression and purification, and enzymatic assays are described in detail in *Supplementary Materials and Methods*.

## RESULTS

### Updated RSH phylogeny across the tree of life

Our previous evolutionary analysis of the RSH protein family applied high-throughput sensitive sequence searching of 1072 genomes from across the tree of life^3^. Since the number of available genomes has grown dramatically in the last decade, we revisited the evolution of RSHs, taking advantage of our new computational tool, FlaGs to analyse the conservation of gene neighbourhoods that might be indicative of functional associations^5^. FlaGs clusters neighbourhood-encoded proteins into homologous groups and outputs a graphical visualisation of the gene neighbourhood and its conservation along with a phylogenetic tree annotated with flanking gene conservation. We identified and classified all the RSHs in 24072 genomes from across the tree of life using our previous Hidden Markov Model (HMM)-based method. We then carried out phylogenetic analysis to identify new subfamilies, generated new HMMs and updated the classification in our database (**Supplementary Tables S1** and **S2**). We have identified 30 subfamilies of SASs, 11 subfamilies of SAHs, and 13 subfamilies of long RSHs (**Figure 1**). The nomenclature follows that of our previous analysis, where prefixes are used to indicate taxonomic distributions^3^.

### Putative toxSAS TA modules are widespread in Actinobacteria, Firmicutes and Proteobacteria

We ran FlaGs on each of all the subfamilies and discovered that Small Alarmone Synthetase (SAS) genes can be frequently found in conserved bicistronic (sometimes overlapping) loci that are characteristic of toxin-antitoxin (TA) loci. Five SAS subfamilies displaying particularly well conserved TA-like arrangements: FaRel (which is actually tricistronic), FaRel2, PhRel, PhRel2 and CapRel (**Figure 2**, **Supplementary File S1** and **Supplementary Table S3**) were selected for further investigation. Among bacteria, PhRel (standing for <u>Ph</u>age Rel, the group to which Gp29^8^ belongs) and FaRel are found in multiple species of <u>F</u>irmicutes and <u>A</u>ctinobacteria (hence the “Fa” prefix), along with representatives of various Proteobacteria; FaRel2 is found in multiple Actinobacteria, and Firmicutes, while PhRel2 is found in firmicutes in addition to bacillus phages. CapRel as a subfamily can be found in a wide diversity of bacteria (including <u>C</u>yanobacteria, <u>A</u>ctinobacteria and <u>P</u>roteobacteria), hence the “Cap” prefix. The putative antitoxins are non-homologous among cognate groups, with the exception of PhRel and CapRel, which share a homologous putative antitoxin (**Figure 2**). PhRel and CapRel are sister groups in the RSH phylogeny with medium support (81% MLB RAxML, 96% UFB IQ-TREE, **Figure 1** and **Supplementary Text S1**), suggesting the TA arrangement has been conserved during the diversification of these groups from a common ancestor.

**Figure 2.**
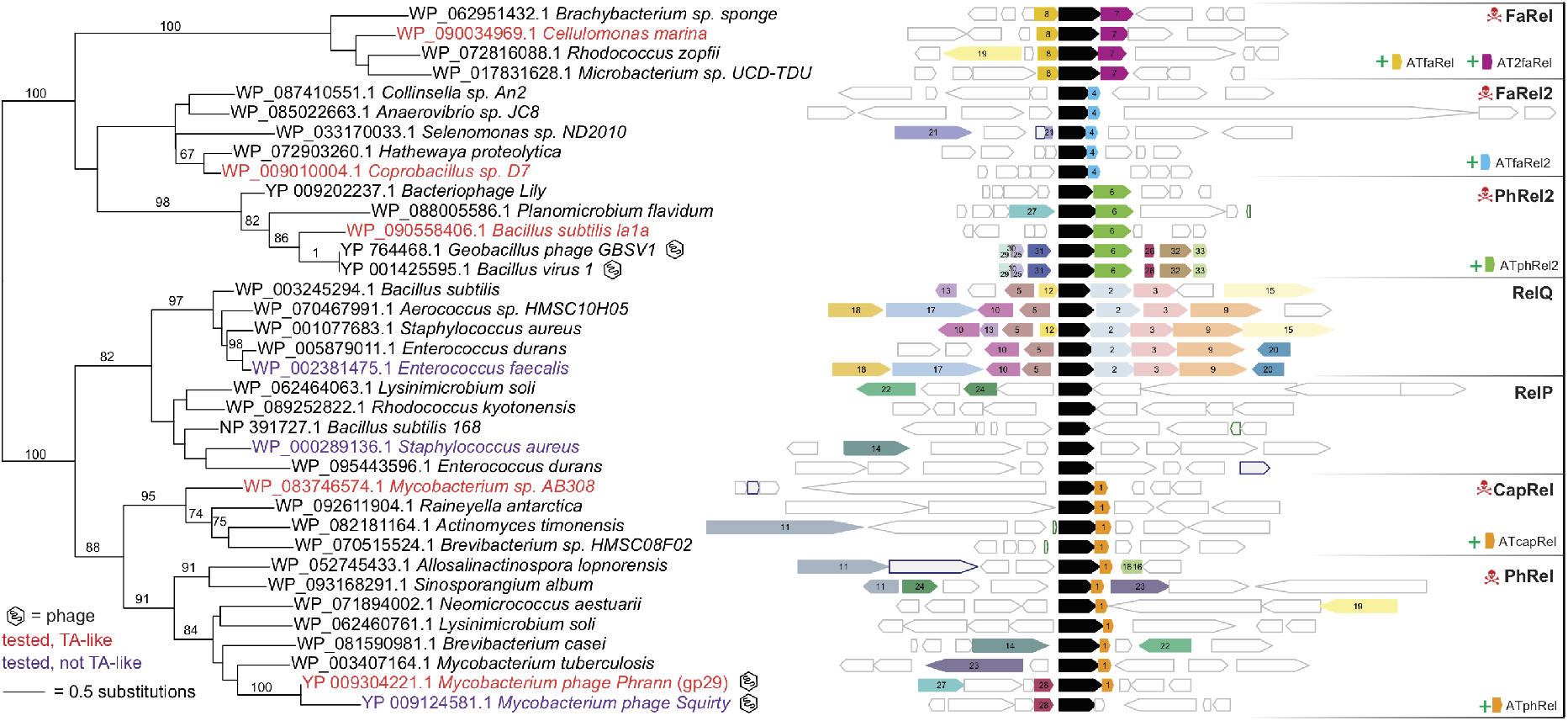
Conservation of gene neighbourhood and maximum likelihood phylogenetic analysis of tested SAS proteins. Genes that encode proteins belonging to a homologous cluster in more than one genomic neighbourhood are coloured and numbered (see **Supplementary Table S3** for the identity of clusters with flanking gene accession numbers). The SAS gene is shown in black, and non-conserved genes are uncoloured. Validated TAs have red taxon names. SASs that we have tested and are non-toxic have purple taxon names. Purple- and green-outlined grey genes are pseudogenes and RNA genes, respectively. Bacteriophage names are indicated with an icon. Numbers on branches are percentage bootstrap support from 100 replicates.

The potential antitoxins are named with an ‘AT’ prefix to the SAS name. ATfaRel is a predicted SAH of the PbcSpo family (**Figure 1**), and ATphRel2 is a GepA (Genetic Element protein A) family homologue. GepA proteins, which carry the DUF4065 domain have previously been associated with TA loci^17^, and are related to the proteolysis-promoting SocA antitoxin of the SocB toxin^18^. The other potential antitoxins (ATcapRel, ATfaRel2, AT2faRel and ATPhRel) have no detectable homology to proteins or domains of known function.

### toxSAS-anti-toxSAS operons encode bona fide type II and type IV TAs

We tested whether SASs encoded in conserved TA-like architectures act as *bona fide* TA systems using a toxicity neutralisation assay in *E. coli* strain BW25113^19^. Putative toxSAS and antitoxin genes were expressed under the control of arabinose- and IPTG-inducible promoters, respectively^19^. Using this approach we have verified five toxSASs as toxic components of *bona fide* TA systems: *Bacillus subtilis* la1a PhRel2 (**Figure 3A**), *Coprobacillus* sp. D7 FaRel2 (**Figure 3B**), *Mycobacterium* phage Phrann PhRel (gp29) (**Figure 3C**) and *Cellulomonas marina* FaRel (**Figure 3D**). Importantly, co-expression of putative antitoxins restored bacterial viability in all of the cases. *C. marina* FaRel is encoded as the central gene in a tricistronic architecture (**Figure 2**), and its toxic effect can be neutralised by expression of either the upstream or, to a lesser extent, the downstream flanking gene (**Figure 3D**). Despite the well-conserved bicistronic organisation, *Mycobacterium tuberculosis* AB308 CapRel (**Figure 3E**) initially displayed no detectable toxicity. Thus, we added a strong Shine-Dalgarno motif (5’-AGGAGG-3’) to increase the translation initiation efficiency in order to drive up its expression levels. In the case of *Mycobacterium sp.* AB308 CapRel, the protein became toxic. Importantly, this toxicity is readily counteracted by the antitoxin ATcapRel (**Figure 3E**). Mycobacterium phage Squirty PhRel^8^ did not display significant toxicity even when the expression was driven with a strong Shine-Dalgarno sequence (**Supplementary Figure S1A**). The reason for this seems to be a large deletion in the synthetase active site in Squirty PhRel (**Supplementary Figure S2**). We also tested well-studied bacterial SASs that are not encoded in TA-like arrangements (*Staphylococcus aureus* RelP^20, 21^ and *Enterococcus faecalis* RelQ^22, 23^). We detected no toxicity, even when the expression is driven by a strong Shine-Dalgarno sequence (**Supplementary Figure S1B**).

**Figure 3.**
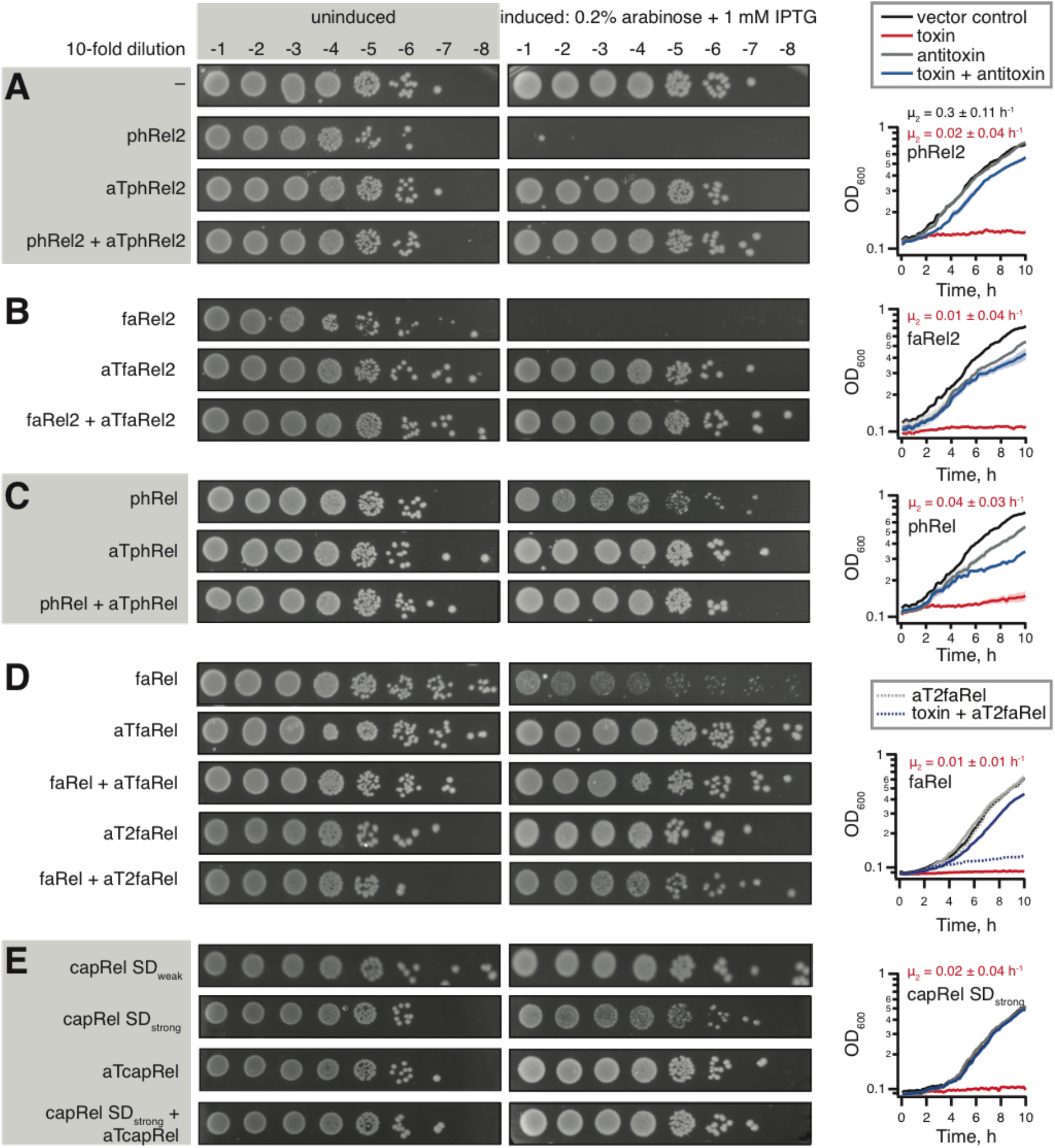
Bi- and tri-cistronic toxSAS operons encode *bona fide* TA systems. Representatives of groups of bi and tri-cistronically encoded toxSAS are validated as TAs: (**A**) *Bacillus subtilis* la1a phRel2:aTphRel2, (**B**) *Coprobacillus* sp. D7 faRel2:aTfaRel2, (**C**) *Mycobacterium* phage Phrann phRel:aTphRel (gp29:gp30) (**D**) *C. marina* faRel:aTfaRel and faRel:aT2faRel and (**E**) *Mycobacterium* sp. AB308 capRel:aTcapRel. To perform the toxicity neutralisation assays on LB plates, overnight cultures of *E. coli* strains transformed with pBAD33 and pKK223-3 vectors or derivatives expressing putative toxSAS toxins and antitoxins, correspondingly, were serially diluted from 10^1^ to 10^8^-fold and spotted on LB medium supplemented with appropriate antibiotics as well as either 1% glucose (repression conditions, left) or 0.2% arabinose and 1 mM IPTG (induction conditions, right). To assay the toxicity in liquid media, bacteria were grown at 37°C in MOPS minimal media supplemented with 0.5% glycerol, 0.2% arabinose and 1mM IPTG (induction conditions). The growth curves represent the geometric mean of three biological replicates and shading represents the standard error, μ_2_ is the growth rate (± standard error) either upon induction of the toxin (in red) or in the absence of the toxin (in black, vector control).

The validated toxSAS toxins differ in the strength of the toxic effect in our system (**Figure 3A-E**): i) FaRel2 and PhRel2 are exceedingly potent and no bacterial growth is detected upon expression of these toxins from the original pBAD33 vector, ii) FaRel and PhRel are significantly weaker and small colonies are readily visible and iii) CapRel is weaker still, with toxicity requiring the introduction of a strong Shine-Dalgarno sequence in the pBAD33 vector. We have validated the observed toxicity by following bacterial growth in liquid culture (**Figure 3**).

Next we tested whether enzymatic activity is responsible for the toxicity of toxSASs. To do so, we substituted a conserved tyrosine in the so-called G-loop for alanine (**Supplementary Figure S3**). This residue is critical for binding the nucleotide substrate and is highly conserved in (p)ppGpp synthetases^24^. All of the tested mutants – PhRel2 Y173A (**Figure 4A**), FaRel2 Y128A, PhRel Y143A and FaRel Y175A – are non-toxic (**Supplementary Figure S3**). Therefore, we conclude that production of a toxic alarmone is, indeed, the universal causative agent of growth inhibition by toxSASs. Finally, the toxicity does not rely on the functionality of the host RSH machinery, since the toxicity phenotype is identical in a Δ*relA* Δ*spoT* (ppGpp^0^) BW25113 *E. coli* strain (**Supplementary Figure S4**).

**Figure 4.**
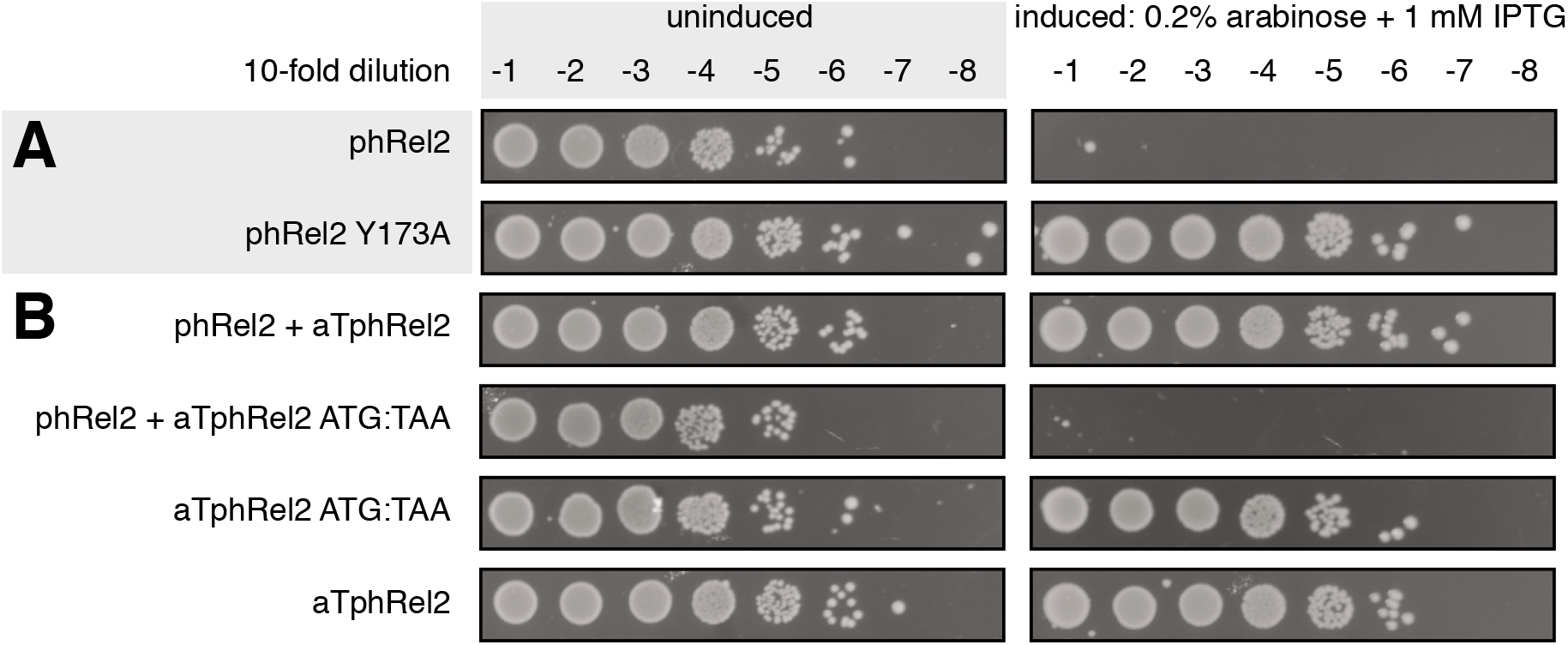
Active site mutations abrogate toxicity of toxSASs, and toxSAS antitoxins work as proteins, not RNA. (**A**) Active site mutation Y173A renders phRel2 toxSAS non-toxic. Analogous experiments with all other identified toxSAS support the essentiality of the enzyme function for toxicity (**Supplementary Figure S3**). (**B**) Mutation of the start codon to stop renders the *aTphRel2* antitoxin ORF unable to protect from the phRel2 toxin. Equivalent experiments of other toxSASs are presented in **Supplementary Figure S5**.

We then investigated whether toxSAS antitoxins inhibit toxSASs on the level of RNA (as in type I and III TA systems) or protein (as in type II and IV TA systems). The former scenario is theoretically possible, since, as we have shown earlier, *E. faecalis* SAS RelQ binds single-stranded RNA and is inhibited in a sequence-specific manner^22^. To discriminate between the two alternatives, we mutated the start codon of the *aTphRel2, aTfaRel2* and *aTphRel* antitoxin ORFs to a stop codon, TAA. Since all of these mutants fail to protect from the cognate toxSAS (**Figure 4B** and **Supplementary Figure S5**), we conclude that they act as proteins, that is, are type II or IV antitoxins.

### The *C. marina* ATfaRel SAH hydrolase antitoxin cross-inhibits all identified toxSAS SASs

The antitoxin ATfaRel is a member of the PbcSpo subfamily of SAH hydrolases (**Figure 1A**). This suggests it acts via degradation of the alarmone nucleotide produced by the toxSAS (and thus as a type IV TA system that does not require direct physical interaction of the TA pair). Therefore, we hypothesised that ATfaRel is able to mitigate the toxicity of all of the identified toxSAS classes through alarmone degradation. This is indeed the case (**Figure 5A**). Similarly, co-expression of human SAH MESH1^25^ universally counteracts the toxicity of toxSASs (**Supplementary Figure S6**). To test if the hydrolysis activity is strictly necessary for antitoxin function, we generated a point mutant of ATfaRel (D54A). Mutation of the homologous active site residue of Rel from *Streptococcus dysgalactiae* subsp. *equisimilis* (Rel_Seq_) abolishes (p)ppGpp hydrolysis^26^. As expected, the D54A mutant is unable to counteract the toxicity from FaRel (**Figure 5B**). The location of the SAS immediately downstream of the SAH raises the question of whether this gene pair has evolved from fission of a long RSH. However, if this was the case, FaRel and ATfaRel would branch in the long RSH part of the phylogeny, which we do not see (**Figure 1**).

**Figure 5.**
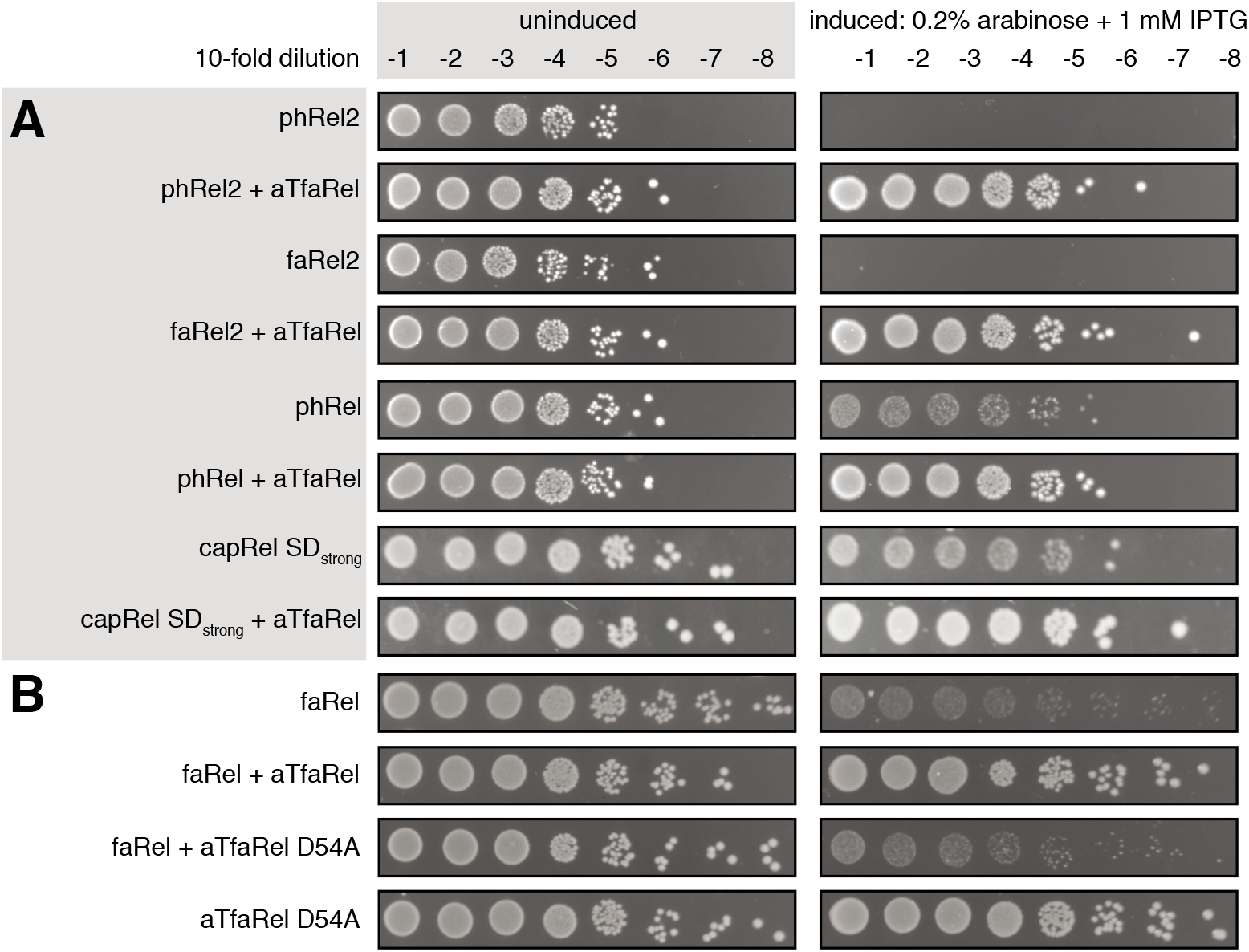
*C. marina* ATfaRel SAH universally counteracts all identified toxSASs. (**A**) *C. marina* aTfaRel neutralises the toxicity of all identified toxSAS toxins. (**B**) Toxicity neutralisation by *C. marina* aTfaRel is abolished by the D54A mutation that inactivates the hydrolytic activity of aTfaRel.

### FaRel toxicity is mediated by accumulation of (p)ppGpp and depletion of ATP and GTP causing rapid inhibition of transcription

To gain first indications for the mechanism of toxSAS-mediated growth inhibition, we assessed the effects of *C. marina* FaRel expression in *E. coli* on overall cell morphology (phase contrast microscopy and FM 5-95 outer membrane staining) and nucleoid appearance (DAPI staining) (**Figure 6A**, **Supplementary Figure S7**). While no change in cell morphology was evident, a rapid decondensation of the nucleoid caused by faRel induction was observed (**Figure 6A** and **Supplementary Figure S8**). This is reminiscent of the decondensation caused by the transcriptional inhibitor rifampicin (**Figure 6A** and **Supplementary Figure S8**), and also by acute RelA-mediated stringent response^27^, suggesting that transcription might be the target of FaRel, To test if *C. marina* FaRel expression indeed inhibits transcription, we assayed macromolecular synthesis rates by following incorporation of ^35^S-methionine in proteins, ^3^H-uridine in RNA and ^3^H-thymidine in DNA (**Figure 6B**, see **Supplementary Figure S9** for method validation). Kanamycin, rifampicin and nalidixic acid were used as controls for specific inhibition of translation, transcription and replication, respectively. Addition of antibiotics causes rapid (within two minutes) inhibition of the corresponding target process (**Figure 6B**, left panel). While expression of *C. marina* FaRel was inhibitory to transcription, translation and replication, the first process to be affected was transcription: the kinetics of inhibition is similar to that of rifampicin (**Figure 6B**, right panel). While the result is in good agreement with (p)ppGpp targeting all of the three processes, the swiftness of the effect on transcription is surprising.

**Figure 6.**
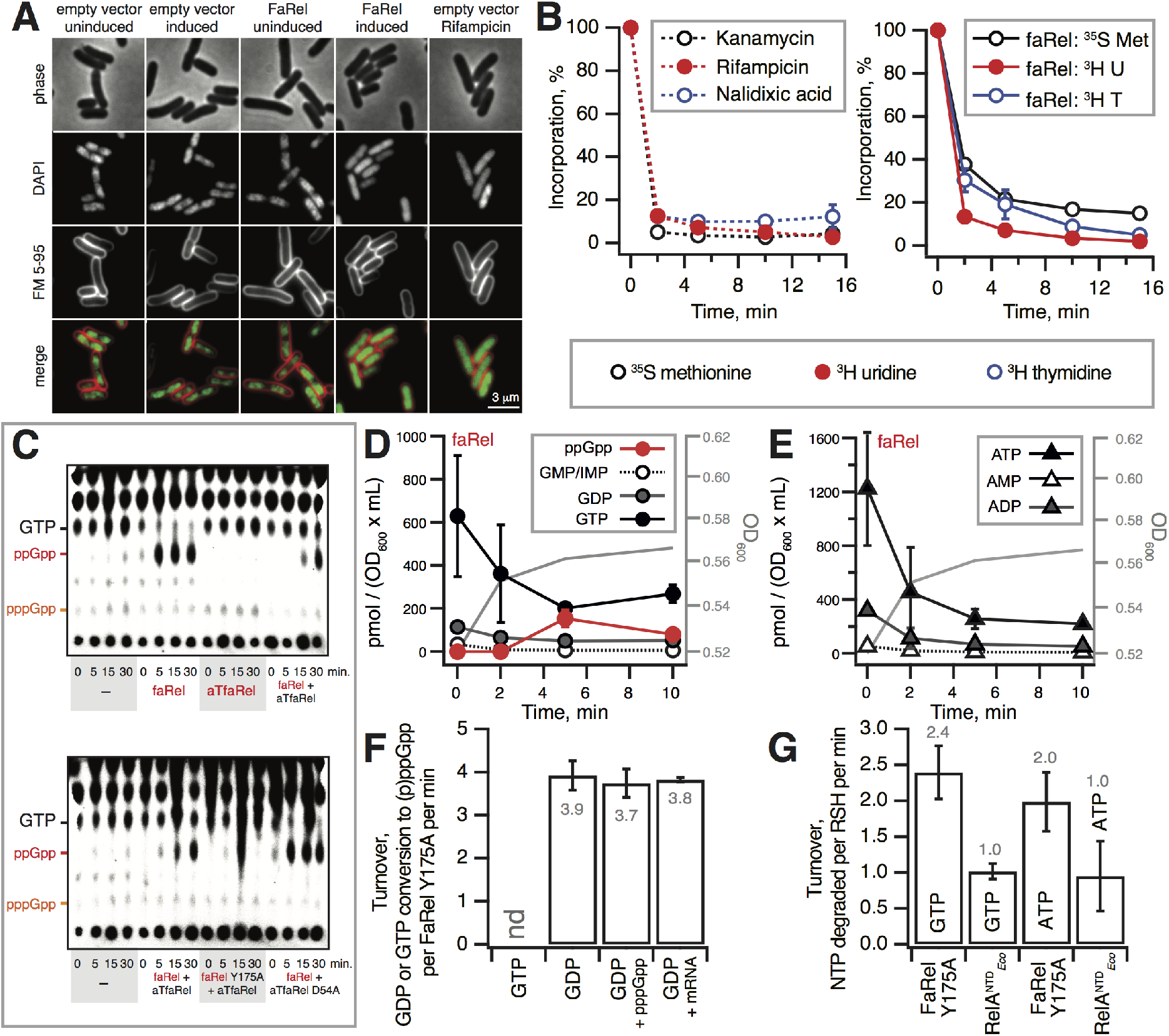
Expression of the *C. marina* FaRel RSH enzyme abrogates transcription by overproducing the (p)ppGpp alarmone and depleting intracellular ATP and GTP. (**A**) Induction of FaRel triggers nucleoid decondensation in *E. coli.* Depicted are phase-contrast (upper panels) and fluorescence images (middle and lower panels) of *E. coli* cells co-stained with DNA-dye DAPI and outer membranedye FM 5-95. The representative cells carry either an empty, or FaRel-expressing vector, and are imaged under uninducing (MOPS-glucose medium) or inducing (15 min in MOPS-glycerol-arabinose medium) conditions. Note the loss of visible nucleoid structure upon induction of FaRel. As a positive control, cells containing empty vector (MOPS-glucose medium) were incubated with rifampicin, which triggers nucleoid decondensation through inhibition of transcription. See **Supplementary Figure S7** for a larger field of view with more cells. (**B**) Pulse-labelling assays following incorporation of ^3^H uridine (black traces), ^35^S methionine (red traces), and ^3^H thymidine (blue traces). *E. coli* BW25113 cells transformed with empty vector control plasmid pBAD33 were treated with 300 μg/ml kanamycin, 100 μg/ml rifampicin and 30 μg/ml nalidixic acid as controls for specific inhibition of translation, transcription and replication, respectively (left panel). Expression of FaRel from the pBAD33-faRel plasmid was induced with 0.2% L-arabinose (right panel). (**C**) The expression of *C. marina* faRel leads to the accumulation of the alarmone ppGpp. Alarmone accumulation is efficiently counteracted by wild type aTfaRel but not its enzymatically compromised D54A mutant. Autoradiograms of a representative TLC plate and a biological replicate (**Supplementary Figure S10**) are presented. (**D** and **E**) Nucleotide pools in *E. coli* BW25113 expressing *C. marina* faRel alone. Cell cultures were grown in defined minimal MOPS medium supplemented with 0.5% glycerol at 37 °C with vigorous aeration. Expression of *C. marina* faRel was induced with 0.2% L-arabinose at the OD_600_ 0.5. Intracellular nucleotides are expressed in pmol per OD_600_ • mL as per the insert. Error bars indicate the standard error of the arithmetic mean of three biological replicates. (**F**) Enzymatic assays with FaRel Y175A in the presence of 300 μM ^3^H GTP or ^3^H GDP combined with 1 mM ATP as substrates, as well as 100 μM pppGpp or 1 μM mRNA(MF). (**G**) Enzymatic assays with *C. marina* faRel Y175A, *E. faecalis* RelQ and *E. coli* RelA^NTD^ in the presence of 1 mM ^3^H ATP or GTP. Experiments were performed in HEPES:Polymix buffer, pH 7.5 at 37 °C in the presence of 5 mM Mg^2+^. Error bars represent SDs of the turnover estimates by linear regression.

We next proceeded to assessing the effects on the intracellular nucleotide pools, with a special focus on (p)ppGpp. First, we used metabolic labelling with ^32^P-orthophosphoric acid combined with TLC separation and autoradiography to assess the accumulation and degradation of nucleotide alarmones upon expression of the *C. marina* FaRel toxSAS and ATfaRel SAH (**Figure 6C** and **Supplementary Figure S10**). The expression of *C. marina* FaRel results in accumulation of ^32^P-ppGpp, which is counteracted by wild type – but not D54A substituted – ATfaRel. While the TLC-based approach is efficient, allowing simultaneous analysis of multiple samples, it lacks the resolution and the quantitative nature of the more laborious HPLC-based approach^28^. Therefore, we analysed the kinetics of nucleotide pools upon expression of either FaRel alone (**Figure 6DE**) or co-expressed with ATfaRel (**Supplementary Figure S11**) by HPLC. Expression of FaRel dramatically perturbs both guanosine (**Figure 5D**) and adenosine (**Figure 5E**) pools. While both GTP and ATP are rapidly depleted, UTP and CTP levels, after the initial drop at two minutes, remain stable (**Supplementary Figure S11**). The result is consistent with neither UTP and CTP serving as substrates for RSH enzymes. The ppGpp levels peak at five minutes and drop at ten. The likely explanation is exhaustion of ATP and GTP that serve as substrates for the RSH enzymes. Efficient depletion of ATP, which is approximately two times more abundant in *E. coli* than GTP (2.2 mM vs 900 μM)^28^ is surprising given that RSH-catalysed pppGpp synthesis is expected to consume guanosines and adenosine in a one-to-one ratio. A possible explanation is that, similarly to a *Streptomyces morookaensis* SAS enzyme^29^, FaRel also catalyses synthesis of pppApp using two ATP molecules as substrates. As judged by our microscopy experiments using the membrane potential-sensitive dye DiSC_3_(5)^30^ and the membrane permeability-indicator Sytox Green^31^, the cells remained both intact and well energised upon expression of FaRel (**Supplementary Figure S8**). Therefore, we can rule out an alternative hypothesis that the reduced nucleotide pools were caused either by FaRel-dependent rapid inhibition of cell metabolism, or by triggered leakage of cytoplasmic content.

The next logical step was to characterise the enzyme biochemically. Despite our best efforts, we failed to express and purify wild type FaRel to homogeneity, even when co-expressed with ATfaRel. We could, however, purify the enzymatically compromised Y175A mutant. Importantly, when overexpressed, FaRel Y175A potently inhibits bacterial growth and this toxicity is counteracted by ATfaRel (**Supplementary Figure S12**), indicating that Y175A and wild type FaRel share the same mechanism of toxicity. We tested the enzymatic activity of FaRel Y175A in the presence of radioactively-labelled ^3^H-GTP or ^3^H-GDP combined with unlabelled ATP (**Figure 6F**). Since *E. faecalis* SAS RelQ is inhibited by single-stranded RNA and activated by pppGpp^22^, we also tested the effects of 1 μM mRNA and 100 μM pppGpp. While we detected no enzymatic activity in the presence of ^3^H-GTP, in the presence of ^3^H-GDP the activity of the catalytically compromised FaRel Y175A is similar to that of wild type *E. faecalis* RelQ^22^. Unlike RelQ – and similarly to *Staphylococcus aureus* RelP^20^ – FaRel Y175A is insensitive to the addition of pppGpp or mRNA(MF) (**Figure 6F**). When FaRel Y175A was incubated with ^3^H-ATP alone, we did not detect ^3^H-pppApp formation. However, both ^3^H-GTP and ^3^H-ATP is degraded, (**Figure 6G**). While we did not detect similar NTP degradation using *E. faecalis* SAS RelQ, the truncated, constitutively active *E. coli* RelA_NTD_ protein also degrades ^3^H-GTP and ^3^H-ATP, though not as efficiently as catalytically compromised FaRel Y175A (**Figure 6G**).

### Class II antitoxins protect only from cognate toxSAS toxins

The gp29-mediated abrogation of growth is employed by the Phrann phage as a defence mechanism against super-infection by other phages^8^. This raises the question of cross-inhibition between toxSAS TA systems: do all of the identified antitoxins inhibit all of the toxSASs (similarly to how the type IV antitoxin SAH ATfaRel protects from all of the tested toxSASs, see **Figure 5A** and **Table 1**) or is the inhibition specific to toxSAS subfamilies TAs? Therefore, we exhaustively tested pairwise combinations of all of the toxSASs with all of the antitoxins (**Table 1** and **Supplementary Figure S13**). ATphRel2, ATfaRel2, ATphRel, ATcapRel and AT2faRel antitoxins could not counteract their non-cognate toxSASs, demonstrating that different classes provide specific discrimination of self from non-self.

**Table 1.** Cross-talk amongst identified toxSAS and their antitoxins as well as standalone phage-encoded SAHs. Toxicity neutralisation assays are presented in (**Figure 5A** and **Supplementary Figures S12** and **S13**). Plus (+) and minus (−) symbols indicate the ability and inability of the antitoxin to neutralise toxicity, respectively.

|  | *Mycobacterium* sp. AB308 ATcapRel | *B. subtilis* la1a ATphRel2 | *Coprobacillus* sp. D7 ATfaRel2 | *Mycobacterium* phage Phrann gp29 ATphRel | *C. marina* ATfaRel SAH | *C. marina* AT2faRel | *Salmonella* phage PVP-SE1 SAH (PbcSpo) | *Salmonella* phage SSU5 SAH (PaSpo) |
| --- | --- | --- | --- | --- | --- | --- | --- | --- |
| CapRel | + | – | – | – | + | – | + | + |
| PhRel2 | – | + | – | – | + | – | + | + |
| FaRel2 | – | – | + | – | + | – | + | + |
| PhRel | – | – | – | + | + | – | + | + |
| FaRel | – | – | – | – | + | + | + | + |

### Numerous SASs and SAHs are encoded in prophage-derived regions of bacterial genomes

Our initial search has identified 13 SASs in bacteriophage genomes, five of which we have confirmed as toxSASs (**Figures 2** and **3**). However, this is likely to be an underestimate for two reasons. Firstly, the currently sequenced phage genomes are a small sample of their entire diversity^32^, and secondly, as prophages reside in bacterial genomes, their genes may not be identified as phage in origin. To detect SAS genes that may be phage in origin but reside in bacterial genomes, we used the tool PHASTER (PHAge Search Tool Enhanced Release^16^), taking a region of DNA equivalent to four upstream and four downstream genes around each SAS and SAH gene (one representative strain per bacterial species). In addition to the already identified phage-encoded CapRel, PhRel and PhRel2, we find 63 prophage regions around representatives in groups belonging to 12 different SASs (**Supplementary Table S4**). It is notable that of RelP and RelQ (the two most broadly distributed SASs), RelP but not RelQ can be phage-associated. An evolutionary history that includes transduction may be part of the reason why the various operon structures of RelP are less well conserved across genera compared with RelQ (**Supplementary File S1**). SAHs are found in many more prophages and prophage-like regions than SASs (90 versus 63 instances, **Supplementary Table S4**). We tested SAHs encoded by Salmonella phages PVP-SE1^33^ (PbcSpo subfamily) and SSU5^34^ (PaSpo subfamily) in toxicity neutralisation assays against validated toxSASs. Like the *C. marina* SAH ATfaRel, both of these stand-alone phage-encoded SAHs efficiently mitigate the toxicity of all the tested toxSASs (**Table 1** and **Supplementary Figure S14**).

## DISCUSSION

Using our tool FlaGs, we have made the surprising discovery that multiple SAS subfamilies can be encoded in TA-like genetic architectures. Through subsequent experimental validation, we have found that the organisation of SAS genes into conserved TA-like bi- (and in one case tri-) cistronic arrangements is an indicator of toxicity. Identification of bicistronic architectures has previously been used as a starting point for prediction of TAs^35, 36^. However, these studies focussed on species that do not encode toxSASs, and therefore these TA systems were not detected. By being associated with novel antitoxins, toxSASs have also escaped identification in “guilt by association” analysis of thousands of genomes^37^. This long-term obscurity is despite toxSAS-containing subfamilies being broadly distributed, present in 239 genera of 15 Gram-positive and -negative phyla of bacterial genomes sampled in this study. Thus, it is likely that there are other previously unknown TA systems to be found that are identifiable through searching for conservation of gene neighbourhoods across disparate lineages, as we have done with FlaGs.

The RSH protein family is widespread; most likely being present in the last common ancestor of bacteria. Thus, for billions of years, these proteins have been used by bacteria to regulate their growth rate in response to their environment by synthesising and hydrolysing nucleotide alarmones. Paradoxically, the very ability of an alarmone to downregulate growth for continued survival is also what gives it toxic potential. We have identified 30 subfamilies of SASs, five of which we have validated as containing toxins, and two of which we have validated as non-toxic (RelP and RelQ). It is likely that SASs exist on a continuum in terms of toxicity, with an antitoxin only being required at a certain level of toxicity. This is supported by the observation that not all toxSASs have the same level of toxicity, with one (*M. tuberculosis* AB308 CapRel) requiring a strong Shine-Dalgarno in order to observe any toxicity in our system. For our five validated toxSAS systems, there are five different homologous groups of antitoxins. This – and the lack of a multi-subfamily toxSAS-specific clade in phylogenetic analysis – suggests toxic SASs could have evolved independently multiple times from non-toxic SASs. In the evolution of a ToxSAS-antiToxSAS module from a non-toxic SAS, it is unlikely that the toxic component evolved before the regulatory antitoxin, as this would be detrimental to fitness. Rather, it is more likely that a SAS became regulated by a neighbouring gene, which relaxed enzymatic constraints on the SAS, allowing it to evolve increased alarmone synthesis rates as well as relax the precision of enzymatic catalysis leading to futile degradation of ATP. While depletion of ATP and GTP pools is expected to contribute to the inhibition of transcription, the fact that the SAH antitoxin efficiently counteracts the toxicity of all ToxSAS SAS enzymes suggests that accumulation of the alarmone is the key toxic effect. We hypothesise that the depletion of the ATP and GTP substrates is responsible for the decrease of (p)ppGpp levels after the initial spike at around five minutes after the induction of FaRel expression. Notably, the (p)ppGpp level remains high in relation to housekeeping ATP and GTP, thus ensuring the efficient shutdown of bacterial growth.

The specific cellular role of most of the toxSASs is unclear, with the exception of the phage PhRel-ATphRel (Gp29-Gp30) toxSAS TA pair, which seems to have a role in inhibition of superinfection^8^. In this system, PhRel encoded by a prophage protects Mycobacteria from infection by a second phage.

Phage infection has previously been linked to alarmone accumulation and stringent response in bacteria^38, 39, 40^. Presumably this is an example of a so-called abortive infection mechanism^41^, where infected hosts are metabolically restricted, but the larger population is protected. A corollary of alarmone-mediated phage inhibition is that incoming phages could bypass this defence system by encoding alarmone hydrolases. Indeed, we have found a variety of different SAHs in different phage genomes and prophage-like regions of bacterial genomes, suggesting there could be cross-talk between ToxSASs and SAHs during infection and superinfection.

## DATA AVAILABILITY

FlaGs is an open source Python application available in the GitHub repository (https://github.com/GCA-VH-lab/FlaGs).

## SUPPLEMENTARY DATA

*Supplementary Data* are available online.

## ACKNOWLEDGMENTS

We are grateful to the Protein Expertise Platform (PEP) Umeå University and Mikael Lindberg for constructing the plasmids used in this work, Mohammad Roghanian for purifying *E. coli* RelA NTD protein, Victoriia Murina for assistance with setting up macromolecular labelling assays, and to Anaïs Poirier for help with toxicity neutralisation experiments.

