## Supplementary_information for "A widespread toxin-antitoxin system exploiting growth control via alarmone signalling": SI_SMethods_SFigures_S1-13_Table_S5_3Sept.pdf

### Supplementary materials

### Supplementary methods

*Construction of plasmids*

All bacterial strains and plasmids used in the study are listed in **Supplementary Table S5**. Oligonucleotides and synthetic genes were synthesized by Eurofins. Toxin ORFs were amplified using primers containing SacI and HindIII restriction sites and cloned in pBAD33 vector. To make the constructs with a strong Shine-Dalgarno motif, the 5'-AGGAGG-3' sequence was incorporated into the pBAD33 vector. The full start codon context including the Shine-Dalgarno motif and intervening sequence was therefore 5'-AGGAGGAATTAAATG-3'. Antitoxin ORFs were amplified using primers containing EcoRI and HindIII restriction sites and cloned in a pKK223-3 vector. Ligation mixes were transformed by heat-shock in *E. coli* DH5$\alpha$. PCR amplifications were carried out using Phusion polymerase, purchased from ThermoScientific along with restriction enzymes and T4 ligase. Point mutations were introduced using QuikChangeKit (Agilent). All final constructs were re-sequenced by Eurofins.

*Microscopy*

Starter cultures were grown overnight at 37 °C in MOPS minimal medium supplemented with 1% glucose, followed by 1:100 dilution into fresh MOPS-glucose medium. The microscope experiments were carried out with cultures grown at 37 °C upon shaking until $OD_{600}$ of 0.3. Induction of the toxin was carried out by washing the cell suspension twice in MOPS medium supplemented with 0.5% glycerol in order to remove glucose, followed by resuspension in the MOPS-glycerol medium additionally supplemented with 0.2% of the inducer arabinose and incubation for 15 min. For maintaining the plasmids, all cultures were grown in the presence of 20 µg/ml chloramphenicol.

When indicated, the cell suspensions were stained for 15 min with 500 ng/ml DAPI (4',6-Diamidino-2-Phenylindole; Sigma-Aldrich), 2 µg/ml FM5-95 (N-(3-Trimethylammoniumpropyl)-4-(6-(4(Diethylamino)phenyl)hexatrienyl) Pyridinium Dibromide; ThermoFisher Scientific), 2 µM $DiSC_3(5)$ (3,3'-Dipropylthiadicarbocyanine Iodine; ThermoFisher Scientific) and 200 nM SYTOX (ThermoFisher Scientific). As positive control for nucleoid decondensation and membrane permeabilisation, respectively, the cell suspensions were incubated for 15 min in the presence of 100 µg/ml Rifampicin or 10 µg/ml Polymyxin B (both Sigma-Aldrich).

The cells were immobilised for microscopy on 1.2% agarose/$H_2O$ as described previously [1]. Microscopy was carried out with Nikon Eclipse Ti equipped with Nikon Plan Apo 100x/1.40 Oil Ph3 objective and Photometrics Prime sCMOS camera. The used light source was either CoolLed pE-300 (CoolLED Ltd.) or Lambda LS (Sutter Instrument). Images were acquired with Metamorph 7.7. The cellular $DiSC_3(5)$ and SYTOX Green fluorescence signals were quantified from background-subtracted micrographs using Fiji [2]. The shown data is representative for three biological replicate experiments.

*Cellular (p)ppGpp quantification by thin layer chromatography (TLC)*

Overnight cultures were pre-grown at 37 °C in liquid MOPS medium supplemented with carbenicillin, chloramphenicol and 1% glucose, diluted to a final $OD_{600}$ of 0.05 in MOPS medium supplemented with

0.5% glycerol and antibiotics, and grown at 37 °C until an $OD_{600}$ of 0.5. At this point the cultures were re-diluted to an $OD_{600}$ of 0.05, and 500 µl aliquots were transferred to 2 ml Eppendorf tubes and spiked with 2.5 µCi of $^{32}$P-orthophosphoric acid (Perkin Elmer). The cultures were grown for two generations (approximately to $OD_{600}$ of 0.2), and expression of the toxins and antitoxins was induced by addition of 0.2% arabinose and 1 mM IPTG (final concentrations), respectively. At 0, 5, 15 and 30 minutes post-induction, 50 µl samples were transferred to 1.5 mL Eppendorf tubes containing 10 µl of 2 M formic acid and pelleted for 2 minutes at 14,000 rpm 4 °C. 10 µl of the resultant supernatant was spotted on PEI Cellulose TLC plates (Merck). The nucleotides were resolved in 1.5 M $KH_2PO_4$, pH 3.4 (VWR Chemicals) and TLC plates were dried and imaged on Phosphorimager Typhoon FLA 9500 (GE Healthcare).

*HPLC-based nucleotide quantification*

*E. coli* strain BW25113[3] was transformed with plasmids pBAD33-faRel and pkk223-3-aTfaRel. The starter cultures were pre-grown overnight at 37 °C in Neidhardt MOPS minimal media[4] supplemented with 1 µg/ml thiamine, 100 µg/ml carbenicillin, 20 µg/ml chloramphenicol and 1% glucose with vigorous shaking (200 rpm). The overnight cultures were diluted to $OD_{600}$ 0.05 in 114 mL of pre-warmed medium MOPS supplemented with 0.5% glycerol as carbon source and grown until $OD_{600} \approx 0.5$ at 37 °C, 200 rpm. At this point combinations of 0.2% arabinose and 1 mM IPTG were added to induce the expression of toxin and antitoxin, respectively. 26 mL samples were collected for HPLC analyses at 0, 2, 5 and 10 minutes after addition of arabinose and IPTG. Nucleotide extraction and HPLC analyses were performed described previously[5]. The $OD_{600}$ measurements were performed in parallel with collecting the samples for HPLC analyses.

*Metabolic labelling with $^{35}$S methionine, $^{3}$H uridine or $^{3}$H thymidine*

One colony of *E. coli* BW25113 cells freshly transformed with either the pBAD33-faRel plasmid for expression of the FaRel toxin or the empty vector control plasmid pBAD33 were used to inoculate overnight cultures in defined Neidhardt MOPS minimal media [4] supplemented with 1% glycerol and appropriate antibiotics. After overnight incubation at 37 °C with shaking, the cultures were diluted to an $OD_{600}$ of 0.05 in 15 mL MOPS minimal media lacking methionine and supplemented with 0.5% glycerol, as well as appropriate antibiotics. The cultures were grown at 37 °C until an $OD_{600}$ of 0.3 in a water bath with shaking (200 rpm), and expression of toxins was induced with 0.2% L-arabinose. Treatment of the cells transformed with pBAD33 vector with 300 µg/ml kanamycin, 100 µg/ml rifampicin and 30 µg/ml nalidixic acid was used with as controls for specific inhibition of translation, transcription and replication, respectively. For a zero point, 1 mL of culture was taken and mixed with either 2 µCi $^{35}$S-methionine (Perkin Elmer), 0.65 µCi $^{3}$H-uridine (Perkin Elmer) or 2 µCi $^{3}$H-thymidine (Perkin Elmer) immediately prior to induction, while simultaneously another 1 mL of culture was taken for $OD_{600}$ measurements. Samples collected 2, 5, 10 and 15 minutes post-induction were treated as described above for the zero point. Incorporation of radioisotopes was quenched after 8 minutes – i.e. on the linear kinetic range thus ensuring that our end-point measurement reflects the rate of incorporation – with addition of 200 µL of ice-cold 50% trichloroacetic acid (TCA). The resultant 1.2 mL culture/TCA samples were loaded onto GF/C filters (Whatman) prewashed with 5% TCA and unincorporated label was removed by washing the filter twice with 5 mL of ice-cold TCA followed by a 5 mL wash with 95% EtOH (twice). The filters were placed in scintillation vials, dried for at least two hours at room temperature, followed by the addition of ScintiSafe 3 scintillation cocktail (5 mL per vial; FisherScientific). After shaking for 15 minutes radioactivity was quantified using TRI-CARB 4910TR 100 V scintillation counter (PerkinElmer).

*Protein expression and purification*

The *C. marina* FaRel Y175A mutant was overexpressed in freshly transformed *E. coli* BW25113 co-transformed with the VHp308 plasmid encoding the aTfaRel antitoxin. Fresh transformants were inoculated to a final $OD_{600}$ of 0.05 in the LB medium (2 L) supplemented with 100 µg/ml carbenicillin, 20 µg/ml chloramphenicol and 0.4 mM IPTG. The cultures were grown at 37 °C until an $OD_{600}$ of 0.4, cooled down for 30 min at 16 °C, induced with 0.2% arabinose (final concentration) and grown for additional 17.5 hours at 16 °C (**Supplementary Figure S12**). The cells were harvested by centrifugation and resuspended in buffer A (750 mM KCl, 500 mM NaCl, 5 mM $MgCl_2$, 40 µM $MnCl_2$, 40 µM $Zn(OAc)_2$, 1 mM mellitic acid, 20 mM imidazole, 10% glycerol, 4 mM β-mercaptoethanol, 25 mM HEPES:KOH pH 8) supplemented with 0.1 mM PMSF and 1 U/ml of DNase I, and lysed by one passage through a high-pressure cell disrupter (Stansted Fluid Power, 150 MPa). Cell debris was removed by centrifugation (25,000 rpm for 1 h) and clarified lysate was taken for protein purification. Mellitic acid was added to buffers as it is known to stabilise the Rel stringent factor from *Thermus thermophilus*[6].

Clarified cell lysate was filtered through a 0.22 µm syringe filter and loaded onto a HisTrap 5 ml HP column pre-equilibrated in buffer A. The column was washed with 5 column volumes (CV) of buffer A, and the protein was eluted with a linear gradient (8 CV, 0-100% buffer B) of buffer B (750 mM KCl, 500 mM NaCl, 5 mM $MgCl_2$, 40 µM $MnCl_2$, 40 µM $Zn(OAc)_2$, 1 mM mellitic acid, 1 M imidazole, 10% glycerol, 4 mM β-mercaptoethanol, 25 mM HEPES:KOH pH 8). Fractions enriched in FaRel Y175A (≈25-50% buffer B) were pooled and concentrated on Amicon Ultra (Millipore) centrifugal filter device (cut-off 10 kDa), totalling approximately 5 ml. The sample was loaded on a HiLoad 16/600 Superdex 200 PG column pre-equilibrated with a high salt buffer (buffer C; 2 M NaCl, 5 mM $MgCl_2$, 10% glycerol, 4 mM β-mercaptoethanol, 25 mM HEPES:KOH pH 8). The fractions containing FaRel Y175A were pooled and applied on a HiPrep 10/26 desalting column (GE Healthcare) pre-equilibrated with storage buffer (buffer D; 720 mM KCl, 5 mM $MgCl_2$, 50 mM arginine, 50 mM glutamic acid, 10% glycerol, 4 mM β-mercaptoethanol, 25 mM HEPES:KOH pH 8). The fractions containing FaRel Y175A were collected and concentrated on an Amicon Ultra (Millipore) centrifugal filter device (cut-off 10 kDa). To cleave off the $His_{10}$-SUMO tag, 83 µg of $His_6$-Ulp1 per 1 mg of FaRel Y175A was added and the reaction mixture was incubated at room temperature for 15 min. After the $His_{10}$-SUMO tag was cleaved off, the protein was passed through a 5 ml HisTrap HP pre-equilibrated with buffer D supplemented with 30 mM imidazole. Fractions containing FaRel Y175A in the flow-through were collected and concentrated on an Amicon Ultra (Millipore) centrifugal filter device with a 10 kDa cut-off. The purity of protein preparations was assessed by SDS-PAGE and spectrophotometrically ($OD_{260}/OD_{280}$ ratio below 0.8 corresponding to less than 5% RNA contamination[7]). Protein preparations were aliquoted, frozen in liquid nitrogen and stored at –80 °C. Individual single-use aliquots were discarded after the experiment.

$His_6$-tagged *E. faecalis* RelQ was overexpressed and purified as described earlier[8]. Untagged *E. coli* $RelA^{NTD}$ (amino acids 1-385) was overexpressed and purified as described earlier for untagged full-length *E. coli* RelA[9].

*Enzymatic assays*

Enzymatic assays were performed as described earlier for *E. faecalis* RelQ[8]., with minor modifications. Experiments were performed in HEPES:Polymix buffer, pH 7.5 at 37 ˚C in the presence of 5 mM $Mg^{2+}$. Error bars represent SDs of the turnover estimates by linear regression.

*$^3$H-ppGpp and $^3$H-pppGpp synthesis assays*: Enzymatic assays were performed with 500 nM FaRel Y175A in the presence of 300 µM $^3$H-GTP or $^3$H-GDP combined with 1 mM ATP as substrates, in the presence or absence of 100 µM pppGpp or 1 µM mRNA(MF).

*$^3$H-GTP and $^3$H-ATP degradation assays*: Enzymatic assays were performed with 500 nM *C. marina* faRel Y175A or *E. faecalis* RelQ or *E. coli* $RelA^{NTD}$ in the presence of 1 mM $^3$H-NTP substrate.

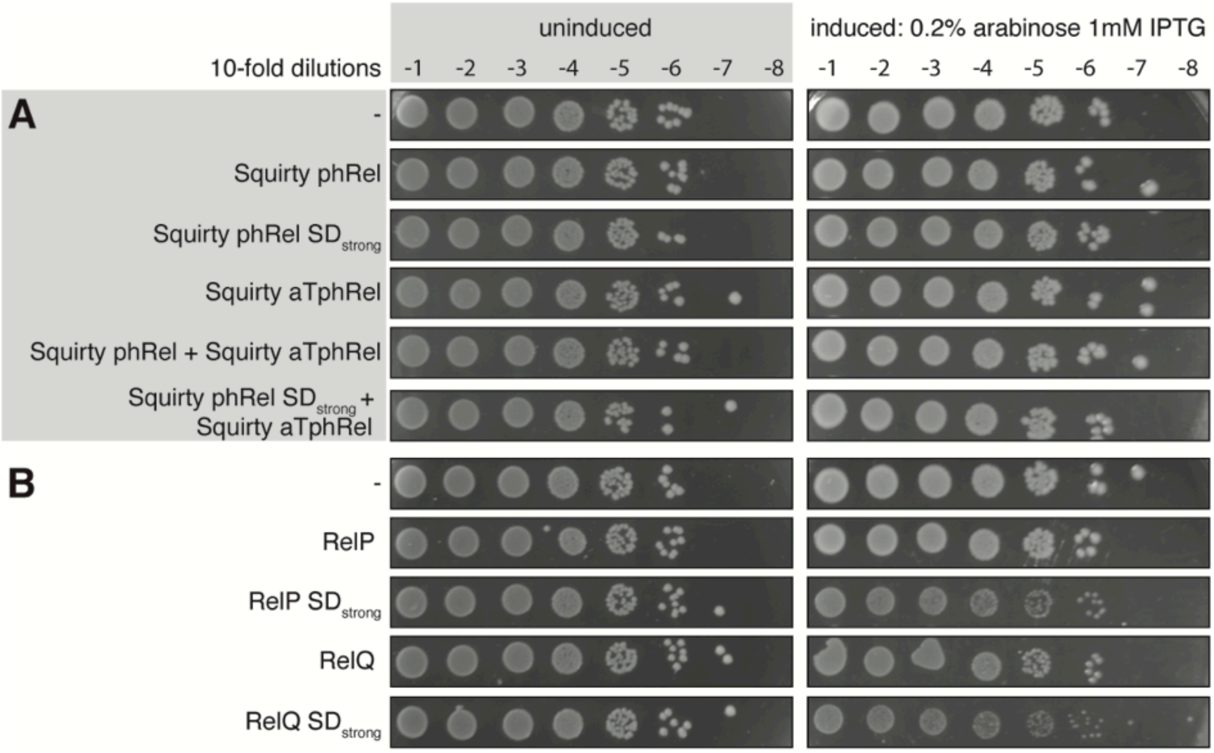

**Supplementary Figure S1 | Toxicity is not a general feature of bacterial SAS enzymes.** Overnight cultures of *E. coli* strains transformed with pBAD33 expressing Squirty phRel, Squirty phRel $SD_{strong}$, *S. aureus* RelP and *E. faecalis* RelQ as well as pKK223-3 expressing Squirty aTphRel or control vectors were serially diluted from $10^1$ to $10^8$-fold and spotted on LB medium supplemented with appropriate antibiotics as well as either 1% glucose (repression conditions, left) or 0.2% arabinose and 1 mM IPTG (induction conditions, right).

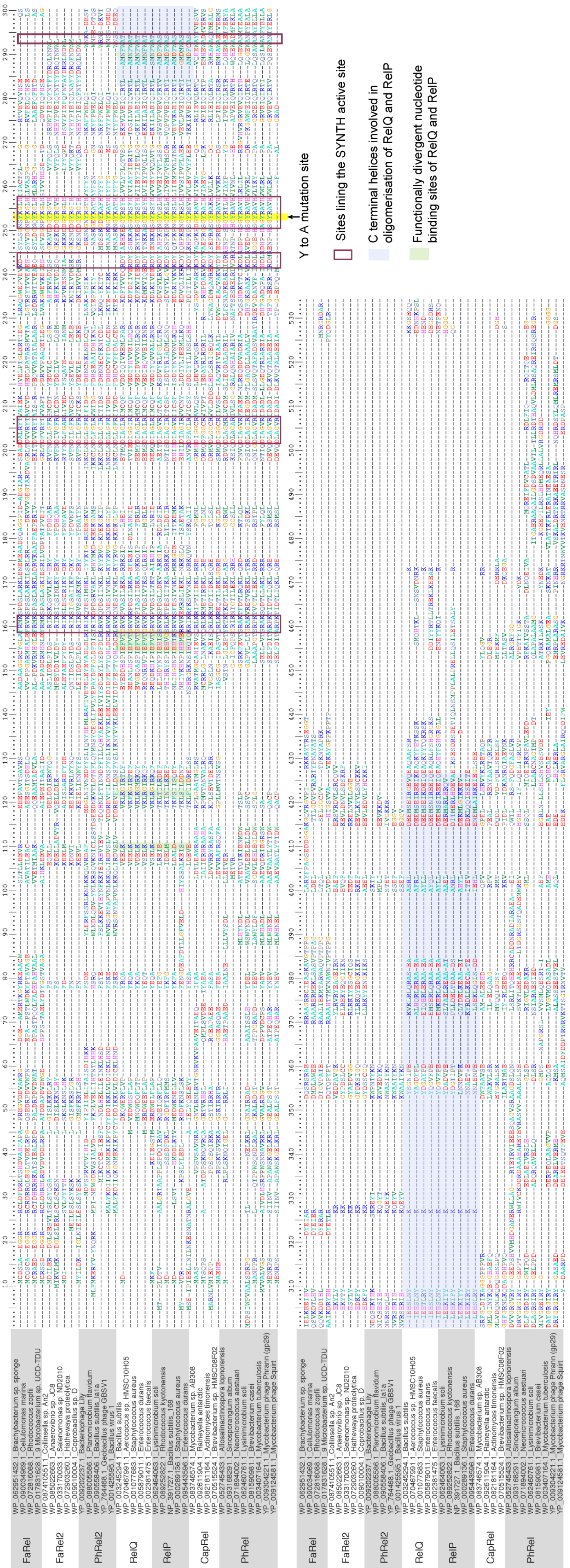

**Supplementary Figure S2 | Alignment of SASs represented in Figure 2, main text.** The alignment is annotated with known functional regions, some of which are specific to RelP and RelQ (*16*). The tyrosine that is substituted for alanine in our mutagenesis analysis is highlighted in yellow.

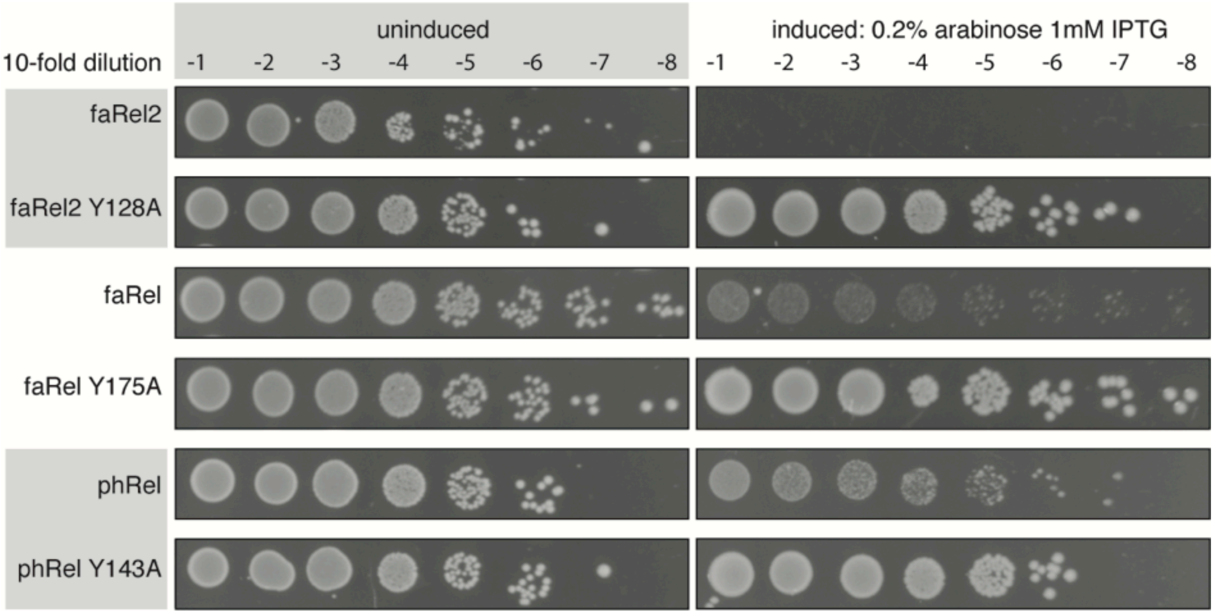

**Supplementary Figure S3 | Substitution of a conserved catalytic Tyrosine residue to Alanine universally abolishes the toxicity of toxSASs.** Overnight cultures of *E. coli* strains transformed with the pBAD33 vector or its derivatives expressing toxSAS toxins (*Coprobacillus* sp. D7 faRel2 wild type or Y128A mutant, *Cellulomonas marina* faRel wild type or Y175A mutant, and *Mycobacterium* phage Phrann phRel (gp29) wild type or Y143A mutant, respectively) were serially diluted from $10^1$ to $10^8$-fold and spotted on LB medium supplemented with appropriate antibiotics as well as either 1% glucose (repression conditions, left) or 0.2% arabinose and 1 mM IPTG (induction conditions, right).

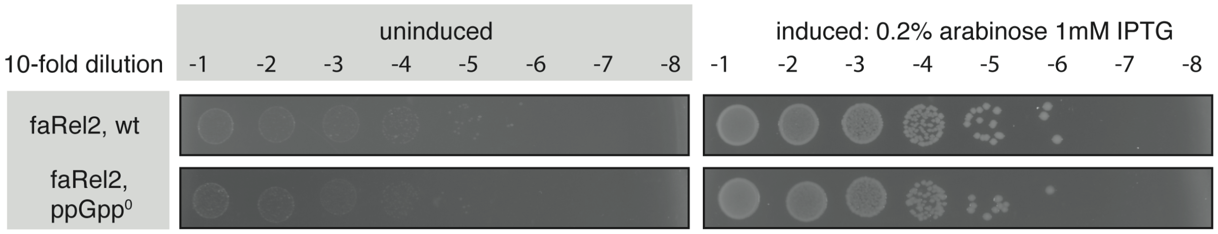

**Supplementary Figure S4 | The toxicity of *Cellulomonas marina* FaRel does not rely on the functionality of the host RSH machinery.** Overnight cultures of *E. coli* Δ*relA* Δ*spoT* (ppGpp$^0$) BW25113 and wild type BW25113 strains transformed with the pBAD33 vector expressing *Cellulomonas marina* faRel were serially diluted from $10^1$ to $10^8$-fold and spotted on LB medium supplemented with appropriate antibiotics as well as either 1% glucose (repression conditions, left) or 0.2% arabinose and 1 mM IPTG (induction conditions, right).

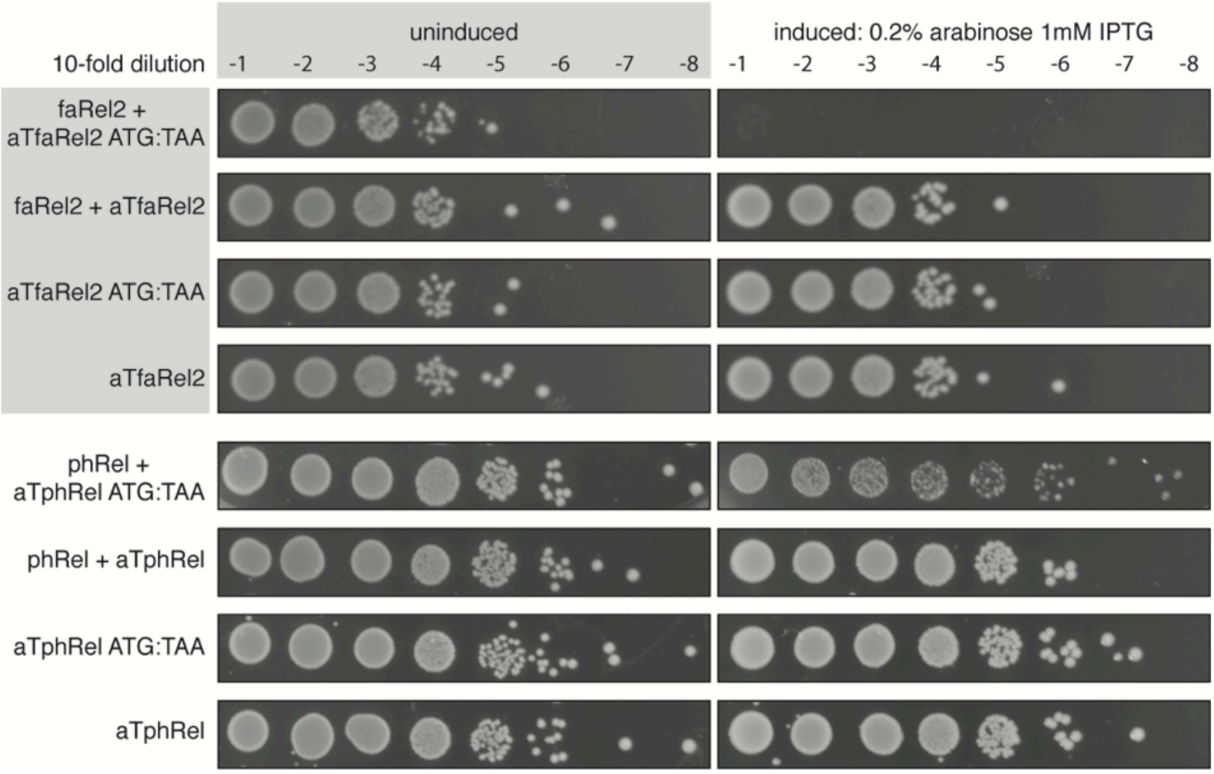

**Supplementary Figure S5 | Replacement of conserved start codon abolishes the ability of the antitoxins to counteract the toxicity of toxSASs.** Overnight cultures of *E. coli* strains transformed with the pBAD33 vector or its derivatives expressing toxSAS toxins (*Coprobacillus* sp. D7 faRel2 and *Mycobacterium* phage Phrann phRel (gp29), respectively) as well as antitoxins (wild type or START-to-STOP mutants) expressed from pKK223-3, were serially diluted from $10^1$ to $10^8$-fold and spotted on LB medium supplemented with appropriate antibiotics as well as either 1% glucose (repression conditions, left) or 0.2% arabinose and 1 mM IPTG (induction conditions, right).

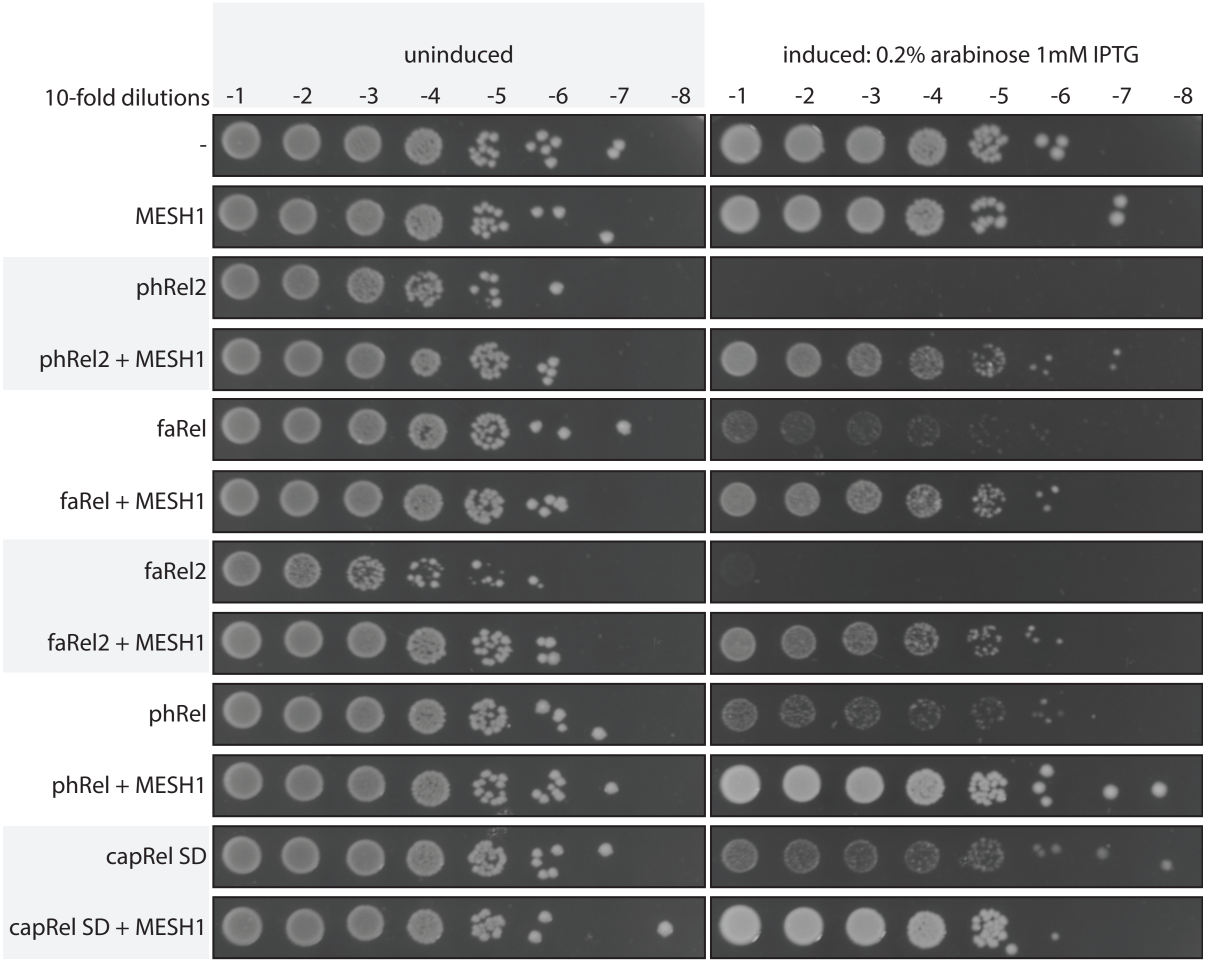

**Supplementary Figure S6 | Human SAH MESH1 universally neutralises the toxicity of toxSASs.** Human MESH1 SAH was tested against all of the identified toxSASs: *B. subtilis* la1a phRel2, *Coprobacillus* sp. D7 faRel2, *C. marina* faRel, *Mycobacterium* phage Phrann phRel (gp29) and *Mycobacterium* sp. AB308 capRel. Overnight cultures of *E. coli* strains transformed with pBAD33 and pKK223-3 vectors or derivatives expressing toxSAS toxins and a phage encoded SAH, respectively, were serially diluted from $10^1$ to $10^8$-fold and spotted on LB medium supplemented with appropriate antibiotics as well as either 1% glucose (repression conditions, left) or 0.2% arabinose and 1 mM IPTG (induction conditions, right).

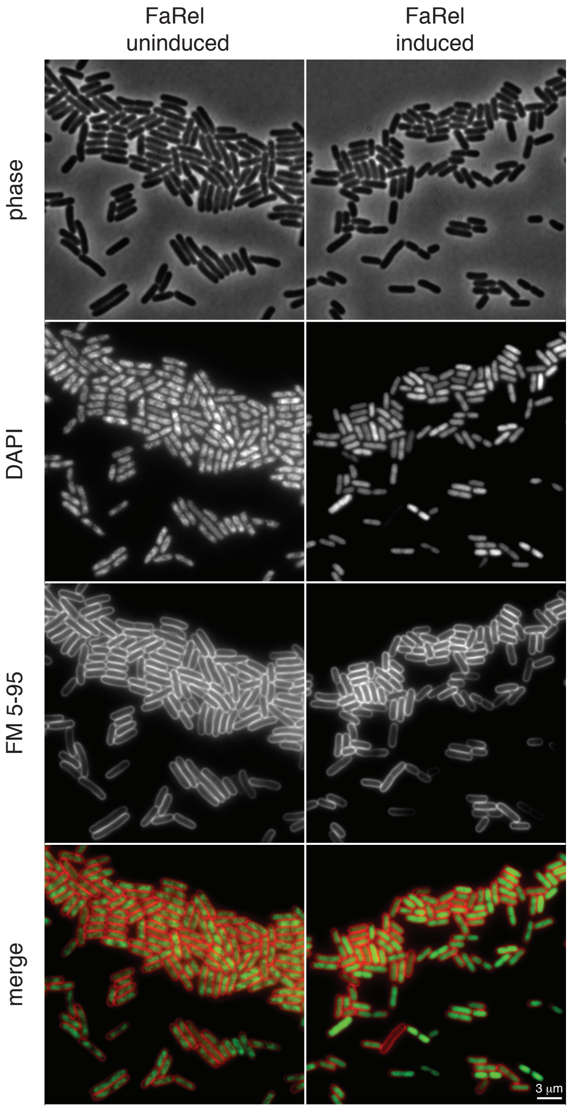

**Supplementary Figure S7 | Induction of FaRel triggers nucleoid decondensation.** Phase-contrast (upper panels) and fluorescence images (middle and lower panels) of *E. coli* cells co-stained with DNA-dye DAPI and membrane-dye FM 5-95. Depicted are representative cells carrying FaRel-expressing vector under uninducing (MOPS-glucose medium) or inducing (15 min in MOPS-glycerol-arabinose medium) conditions. Note the loss of visible nucleoid structure upon induction of FaRel. This a larger field of view of cells shown in **Figure 6**.

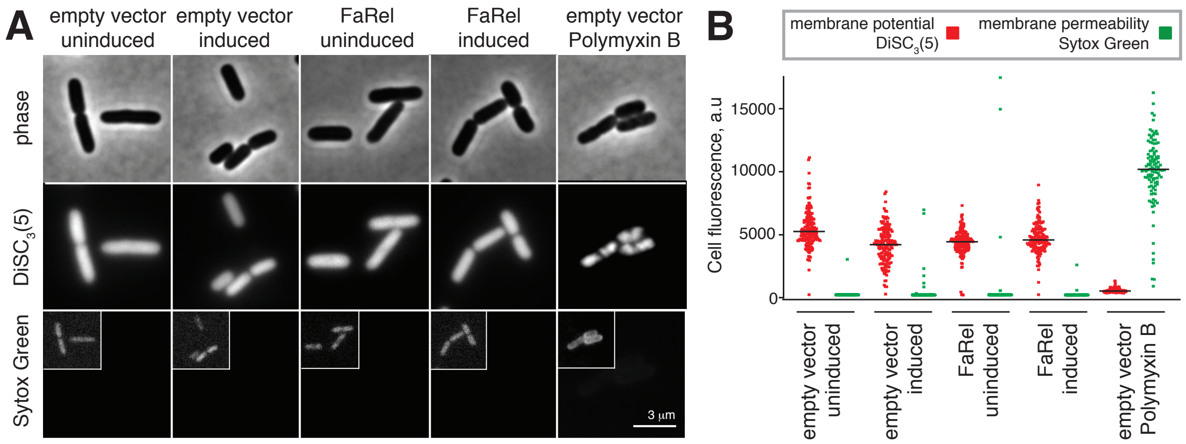

**Supplementary Figure S8 | Induction of FaRel does not interfere with membrane integrity or oxidative phosphorylation.** (**A**) Phase-contrast (upper panels) and fluorescence images (middle and lower panels) of *E. coli* cells co-stained with membrane potential-sensitive dye $DiSC_3(5)$ and membrane permeability-indicator Sytox Green, respectively. Depicted are representative cells carrying either an empty, or FaRel-expressing vector under uninducing (MOPS-glucose medium) or inducing (15 min in MOPS-glycerol-arabinose medium) conditions. As a positive control, cells containing empty vector (MOPS-glucose medium) were incubated for 15 min with membrane depolarising and permeabilising antibiotic Polymyxin B. High $DiSC_3(5)$-fluorescence levels indicate high membrane potential levels found in well energised, metabolically active cells. High Sytox Green-levels, in contrast, indicate cells with permeabilised cytoplasmic membranes. The large depicted images preserve the fluorescence intensity differences between the individual conditions. The small insert images serve to visualise the presence of cells in otherwise dark image fields. (**B**) Fluorescence quantification for a larger number of cells from the same imaging dataset (n = 115-184 cells).

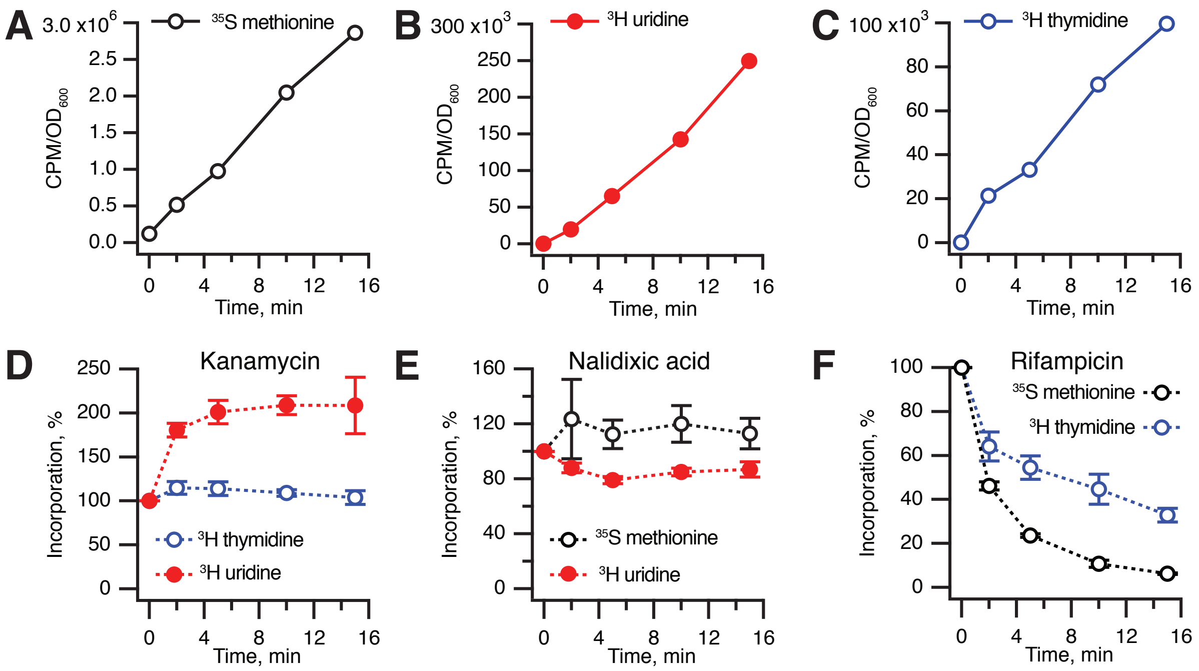

**Supplementary Figure S9 | Control experiments for pulse-labelling assays.** Pulse-labelling assays following kinetics of incorporation of $^{3}$H uridine (**A**, black traces), $^{35}$S methionine (**B**, red traces), and $^{3}$H thymidine (**C**, blue traces**)** in *E. coli* BW25113 ($OD_{600}$ 0.5) growing in MOPS-glucose medium at 37 °C. Kinetics of $^{3}$H uridine, $^{35}$S methionine and $^{3}$H thymidine incorporation in *E. coli* BW25113 ($OD_{600}$ 0.5) upon addition of with 300 µg/ml kanamycin (**D**), 30 µg/ml nalidixic acid (**E**) or 100 µg/ml rifampicin (**F**). The efficiency of incorporation is normalised to untreated control. Error bars indicate the standard error of the arithmetic mean of three biological replicates.

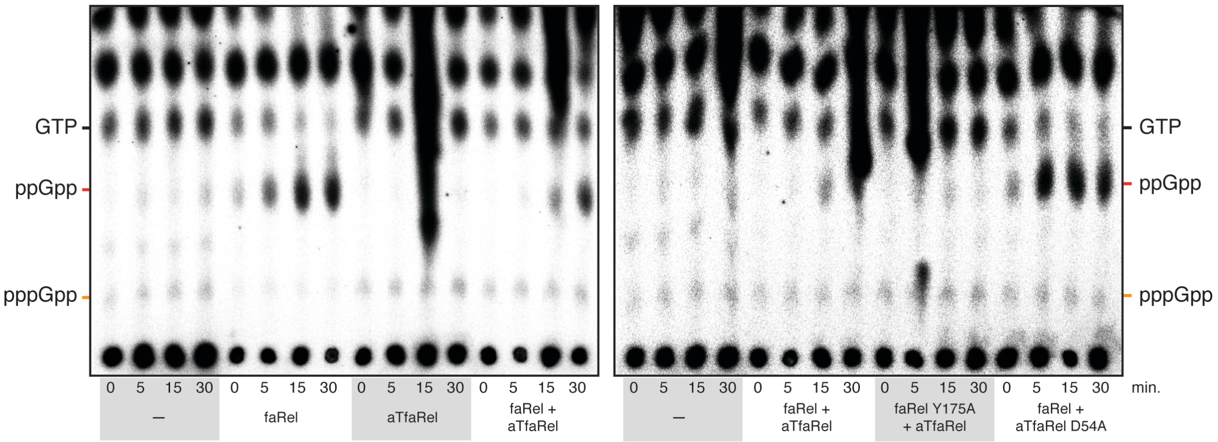

**Supplementary Figure S10 | The *C. marina* ATfaRel SAH efficiently degrades (p)ppGpp produced by the FaRel toxSAS in live *E. coli* cells.** The expression of *C. marina* faRel leads to the accumulation of the alarmone ppGpp which is efficiently counteracted by wild type aTfaRel but not its enzymatically-compromised D54A mutant. Autoradiograms of a biological replicate are shown.

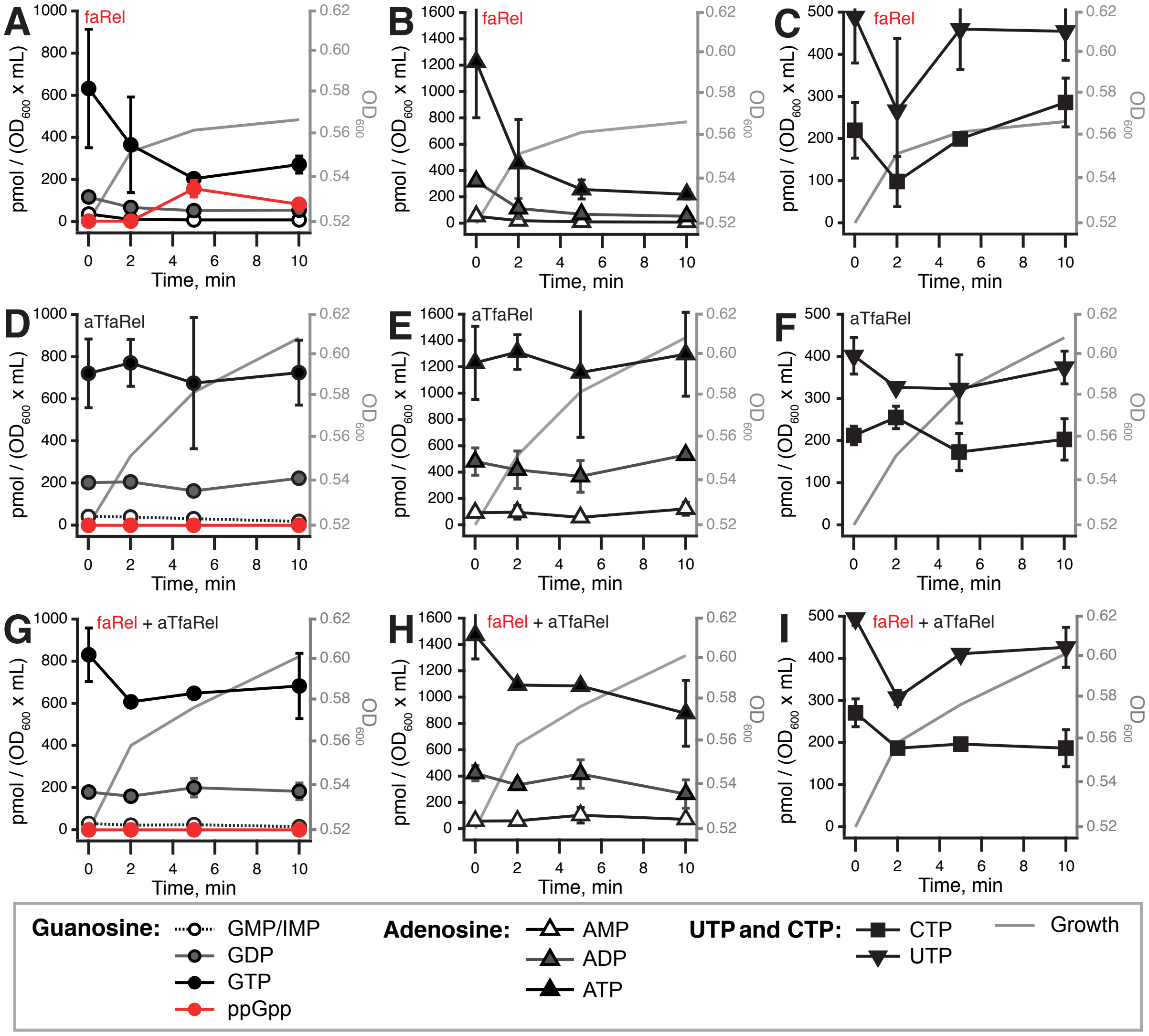

**Supplementary Figure S11 | Nucleotide pools in *E. coli* BW25113 expressing *C. marina* faRel (A-C), *C. marina* aTfaRel (D-F), and the combination of *C. marina* faRel and aTfaRel (G-I).** Cell cultures were grown in defined minimal MOPS medium supplemented with 0.5% glycerol at 37 °C with vigorous aeration. The expression of *C. marina* faRel was induced with 0.2% L-arabinose at the $OD_{600}$ 0.5 (**A-C**, **G-I**). The expression of *C. marina* aTfaRel was induced by 1 mM IPTG at the zero time point (**D-F**, **G-I**). Intracellular nucleotides are expressed in pmol per $OD_{600}$ • mL as per the insert. Error bars indicate the standard error of the arithmetic mean of three biological replicates.

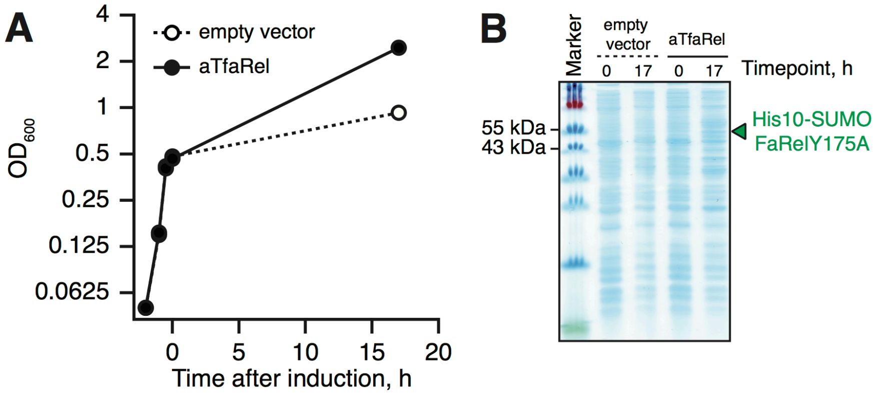

**Supplementary Figure S12 | Growth inhibition mediated by overexpression of the *C. marina* faRel Y175A mutant is counteracted by *C. marina* aTfaRel.** *E. coli* cultures were grown in LB medium supplemented with 0.4 mM IPTG at 37 °C until an $OD_{600}$ of 0.4, shifted to 16 °C and expression of *C. marina* faRel Y175A was induced by addition of arabinose to a final concentration of 0.2%.

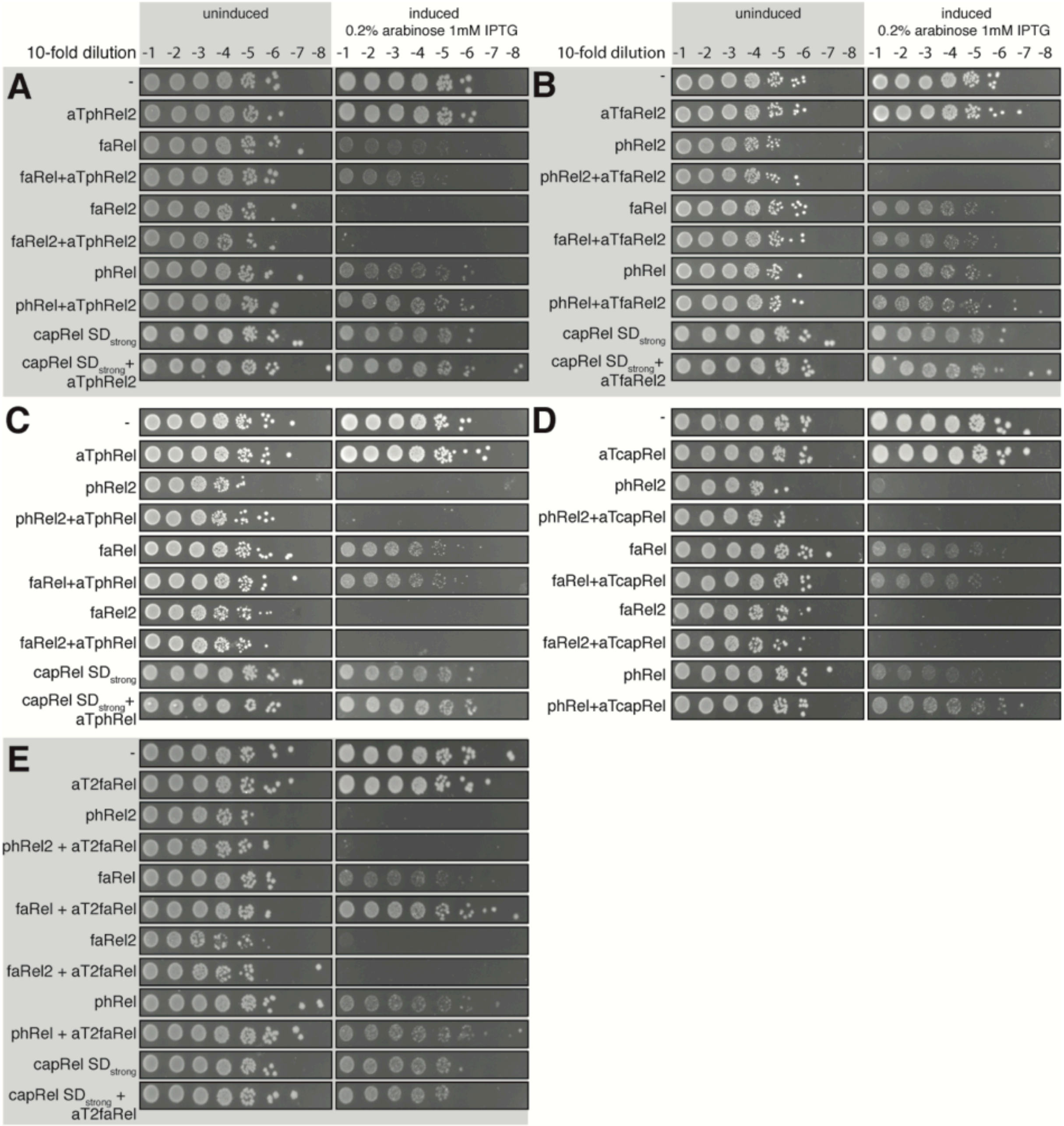

**Supplementary Figure S13 | Cross-reactivity among toxSAS toxins and antitoxins.** Overnight cultures of *E. coli* strains transformed with pBAD33 vector or its derivatives expressing toxSAS toxins (*B. subtilis* la1a phRel2, *Coprobacillus* sp. D7 faRel2, *C. marina* faRel, *Mycobacterium* phage Phrann phRel (gp29), and *Mycobacterium* AB308 capRel), as well as antitoxins aTphRel2 (**A**), aTfaRel2 (**B**), aTphRel (**C**), aTcapRel (**D**) and aT2faRel (**E**) expressed from pKK223-3, were serially diluted from $10^1$ to $10^8$-fold and spotted on LB medium supplemented with appropriate antibiotics as well as either 1% glucose (repression conditions, left) or 0.2% arabinose and 1 mM IPTG (induction conditions, right).

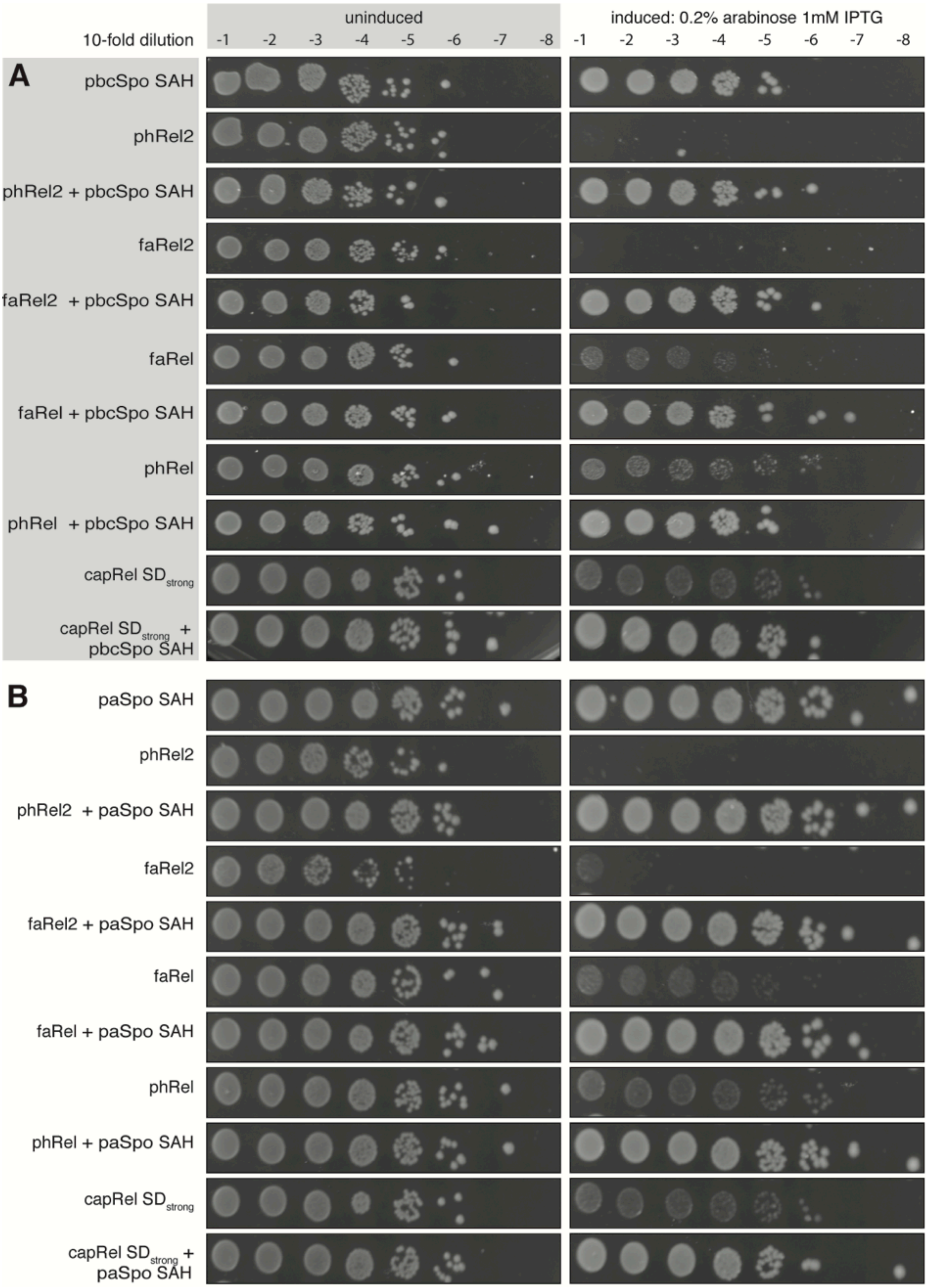

**Supplementary Figure S14 | The standalone SAH encoded by *Salmonella* phages PVP-SE1 and SSU5 universally neutralise the toxicity of toxSAS.** pbcSpo SAH encoded by *Salmonella* phage PVP (**A**) and paSpo SAH encoded by *Salmonella* phage SSU5 (**B**) tested against all of the identified toxSASs: *B. subtilis* la1a phRel2, *Coprobacillus* sp. D7 faRel2, *C. marina* faRel, *Mycobacterium* phage Phrann phRel (gp29) and *Mycobacterium* sp. AB308 capRel. Overnight cultures of *E. coli* strains transformed with pBAD33 and pKK223-3 vectors or derivatives expressing toxSAS toxins and a phage encoded SAH, respectively, were serially diluted from $10^1$ to $10^8$-fold and spotted on LB medium supplemented with appropriate antibiotics as well as either 1% glucose (repression conditions, left) or 0.2% arabinose and 1 mM IPTG (induction conditions, right).

**Supplementary Table S5. Strains and plasmids used in this study**

| Annotation | Strains | Description | Reference/source |
|---|---|---|---|
| VHb 17 | *E. coli* BW25113 F-, Δ(*araD-araB*)*567*, Δ*lacZ4787*(::rrnB-3), *λ*-, *rph-1*, Δ(*rhaD-rha*B)*568*, *hsdR514* | Wild type *E. coli* BW25113 | [3] |
| VHb 283 | *E. coli* BW25113 Δ*relA* Δ*spoT* Δ(*araD-araB*)*567*, Δ*lacZ4787*(::rrnB-3), *λ*-, *rph-1*, Δ(*rhaD-rha*B)*568*, *hsdR514* | ppGpp$^0$ *E. coli* BW25113 | [10] |
| | **Plasmids** | | |
| VHp 83 | pET21a:RelQEf | Expression construct for C-terminally His$_6$-tagged *E. faecalis* RelQ SAS | [11] |
| VHp173 | pET24d::*his$_{10}$-SUMO-relA$^{NTD}$* | Expression construct for C-terminally His$_{10}$-SUMO-tagged *E. coli* RelA$^{NTD}$ (residues 1-385) | [9] |
| VHp 213 | pBAD33 | P15A, Cml$^R$, pBAD promoter | [12] |
| VHp 214 | pkk223-3 | ColE1, Amp$^R$, pTac promoter | [13] |
| VHp 220 | pBAD33-phRel (Phrann) | phRel toxin from *Mycobacterium* Phage Phrann under control of pBAD promoter | This work |
| VHp 221 | pkk223-3-aTphRel (Phrann) | aTphRel antitoxin from *Mycobacterium* Phage Phrann under control of pTac promoter | This work |
| VHp 254 | pBAD33-phRel Y143A | phRel toxin with Y143A mutation under control of pBAD promoter | This work |
| VHp 277 | pBAD33-faRel2 | faRel2 toxin under control of pBAD promoter | This work |
| VHp 278 | pkk223-3-aTfaRel2 | aTfaRel2 antitoxin under control of pTac promoter | This work |
| VHp 279 | pBAD33-phRel (Squirty) | phRel toxin from *Mycobacterium* Phage Squirty under control of pBAD promoter | This work |
| VHp 280 | pkk223-3-aTphRel (Squirty) | aTphRel antitoxin from *Mycobacterium* Phage Squirty under | This work |

| | | | |
|---|---|---|---|
| | | control of pTac promoter | |
| VHp 303 | pBAD33-phRel2 | phRel2 toxin under control of pBAD promoter | This work |
| VHp 304 | pkk223-3-aTphRel2 | aTphRel2 antitoxin under control of pTac promoter | This work |
| VHp 307 | pBAD33-faRel | faRel toxin under control of pBAD promoter | This work |
| VHp 308 | pkk223-3-aTfaRel | aTfaRel antitoxin under control of pTac promoter | This work |
| VHp 311 | pBAD33-S. aureus RelP | *Staphylococcus aureus* RelP under control of pBAD promoter | This work |
| VHp 312 | pBAD33-E. faecalis RelQ | *Enterococcus faecalis* RelQ under control of pBAD promoter | This work |
| VHp 331 | pkk223-3-aTfaRel2 ATG:TAA | aTfaRel2 antitoxin start:stop codon mutation under control of pTac promoter | This work |
| VHp 335 | pBAD33-capRel | capRel toxin under control of pBAD promoter | This work |
| VHp 336 | pkk223-3-aTcapRel | aTcapRel antitoxin under control of pTac promoter | This work |
| VHp 358 | pkk223-3-aTfaRel D54A | aTfaRel antitoxin with D54A mutation under control of pTac promoter | This work |
| VHp 360 | pBAD33-phRel2 Y173A | phRel2 toxin with Y173A mutation under control of pBAD promoter | This work |
| VHp 361 | pBAD33-faRel Y175A | faRel toxin with Y175A mutation under control of pBAD promoter | This work |
| VHp 366 | pBAD33-faRel2 Y128A | faRel2 toxin with Y128A mutation under control of pBAD promoter | This work |
| VHp 367 | pkk223-3-aTphRel ATG:TAA | aTphRel with a start:stop codon mutation under control of pTac promoter | This work |
| VHp 379 | pkk223-3-aTphRel2 ATG:TAA | aTphRel2 antitoxin with a start:stop codon mutation under control of pTac promoter | This work |
| VHp 380 | pBAD33-capRel $SD_{strong}$ | capRel toxin with a strong Shine-Dalgarno sequence under control of pBAD promoter | This work |
| VHp 427 | pKK223-3-hMESH1 | Human MESH1 SAH under control of pTac promoter | This work |
| VHp 484 | pBAD33-His10-SUMO- | N-terminally $His_{10}$-SUMO-tagged | This work |

| | | | |
|---|---|---|---|
| | faRel Y175A $SD_{strong}$ | faRel toxin with Y175A mutation under control of pBAD promoter. strong Shine-Dalgarno sequence | |
| VHp 509 | pKK223-3-aT2faRel | aT2faRel antitoxin under control of pTac promoter | This work |
| VHp 510 | pBAD33-phRel (Squirty) $SD_{strong}$ | phRel toxin with a strong Shine-Dalgarno sequence from *Mycobacterium* Phage Squirty under control of pBAD promoter | This work |
| VHp 511 | pBAD33-S. aureus RelP $SD_{strong}$ | *Staphylococcus aureus* RelP with a strong Shine-Dalgarno sequence under control of pBAD promoter | This work |
| VHp512 | pBAD33-E. faecalis RelQ $SD_{strong}$ | *Enterococcus faecalis* RelQ with a strong Shine-Dalgarno sequence under control of pBAD promoter | This work |

**Supplementary legends for additional SI files:**

**Supplementary Table S1. Classification of 35615 RSH sequences into subfamilies, accompanied by domain assignments**
The unique sequence identifiers are included for retrieval of data from online repositories.

**Supplementary Table S2. Taxonomy of 24072 species, and their RSH composition**
All species considered in the analysis are listed, ordered by taxonomy. The number and identity of RSH subfamilies are recorded.

**Supplementary Table S3. Homologous cluster legends for FlaGs output shown in Figure 2 and Supplementary File S1, showing the protein accessions and titles of each protein in each numbered cluster.**
Data sets are one per sheet.

**Supplementary Table S4. PHASTER results showing the phage sequences found in protein accession from different SAH and SAS subfamilies.**
Data sets from SAHs and SASs are one per sheet.

**Supplementary File S1. Conservation of neighbourhood around ToxSAS genes.**
This is the FlaGs graphical output for the SAS subfamilies (one representative strain per species for toxSASs and one representative per genus for RelP and RelQ) that contain the 5 toxSASs, in PDF format. Genes that encode proteins belonging to a homologous group in more than one genomic neighbourhood are coloured and numbered. The SAS gene is shown in black, and uncoloured genes are unconserved. The legend showing protein content of clusters, with accession numbers and descriptions, is available in **Supplementary Table S3**.

**Supplementary Text S1. The sequence alignments used to generate Figure 1.**
All alignments are in the same file, in FASTA format, separated by comments preceded by “#”.
