## Supplementary figures and images for "A widespread toxin-antitoxin system exploiting growth control via alarmone signalling"

### Supplementary_File_S1.pdf

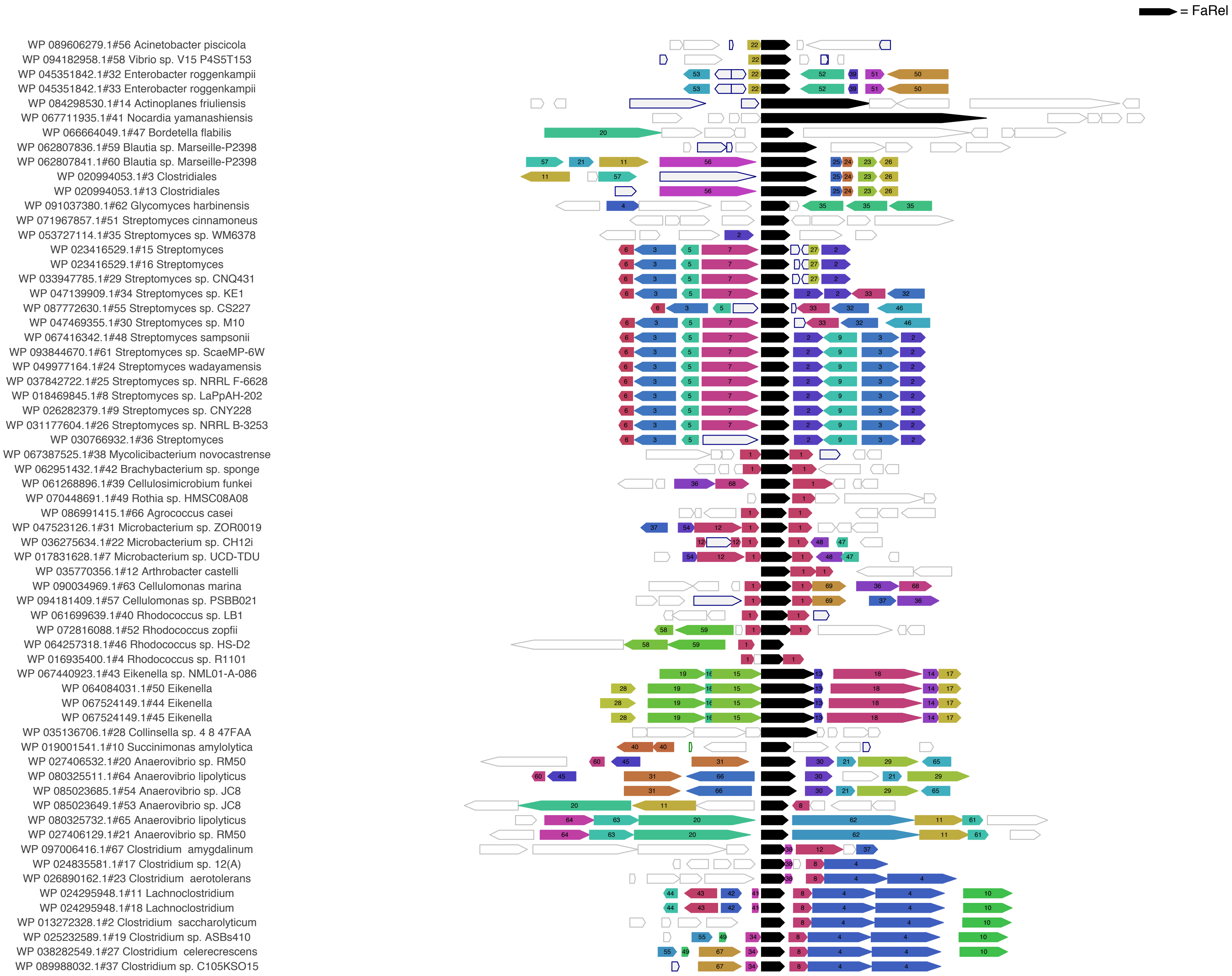

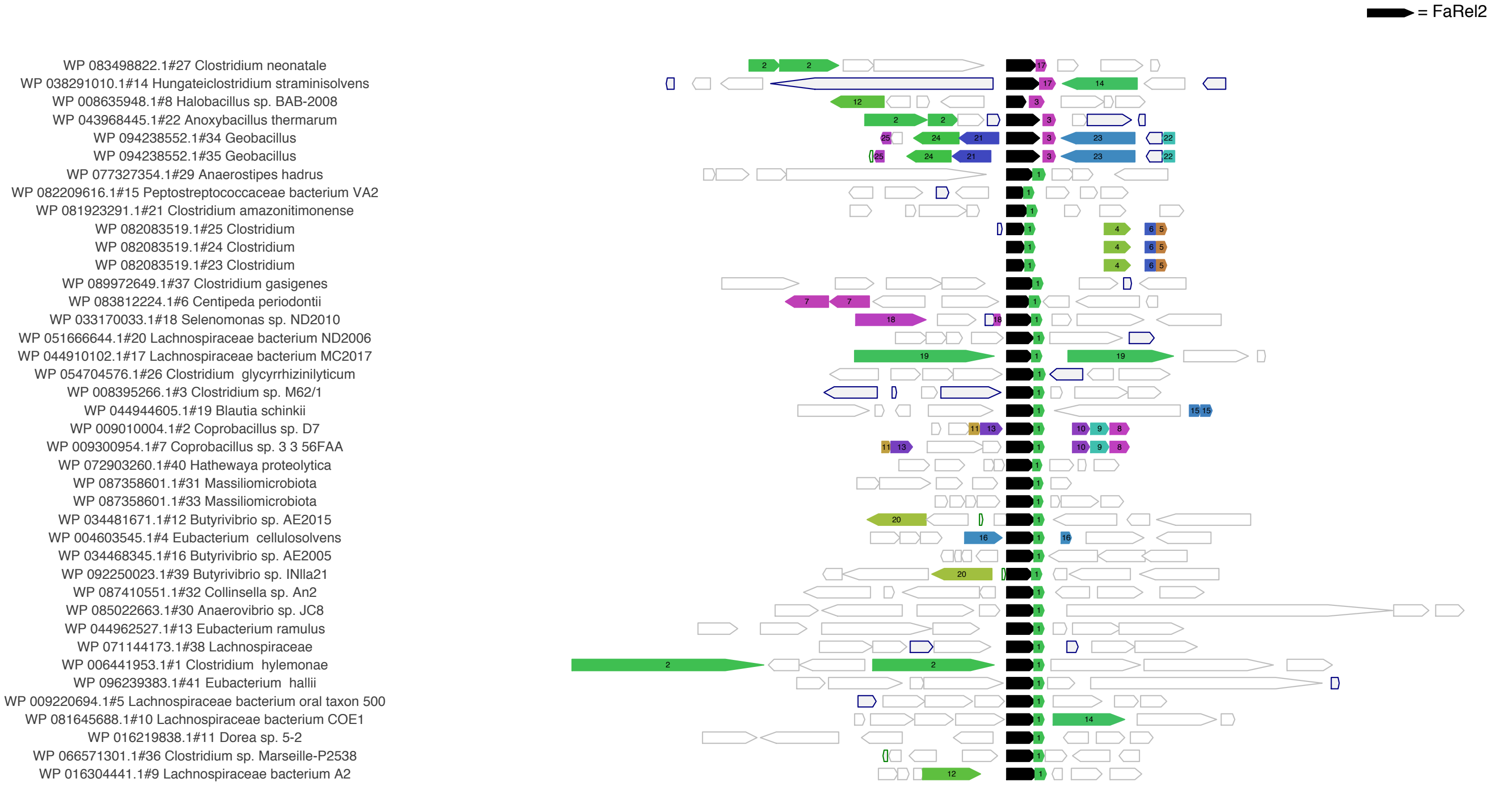

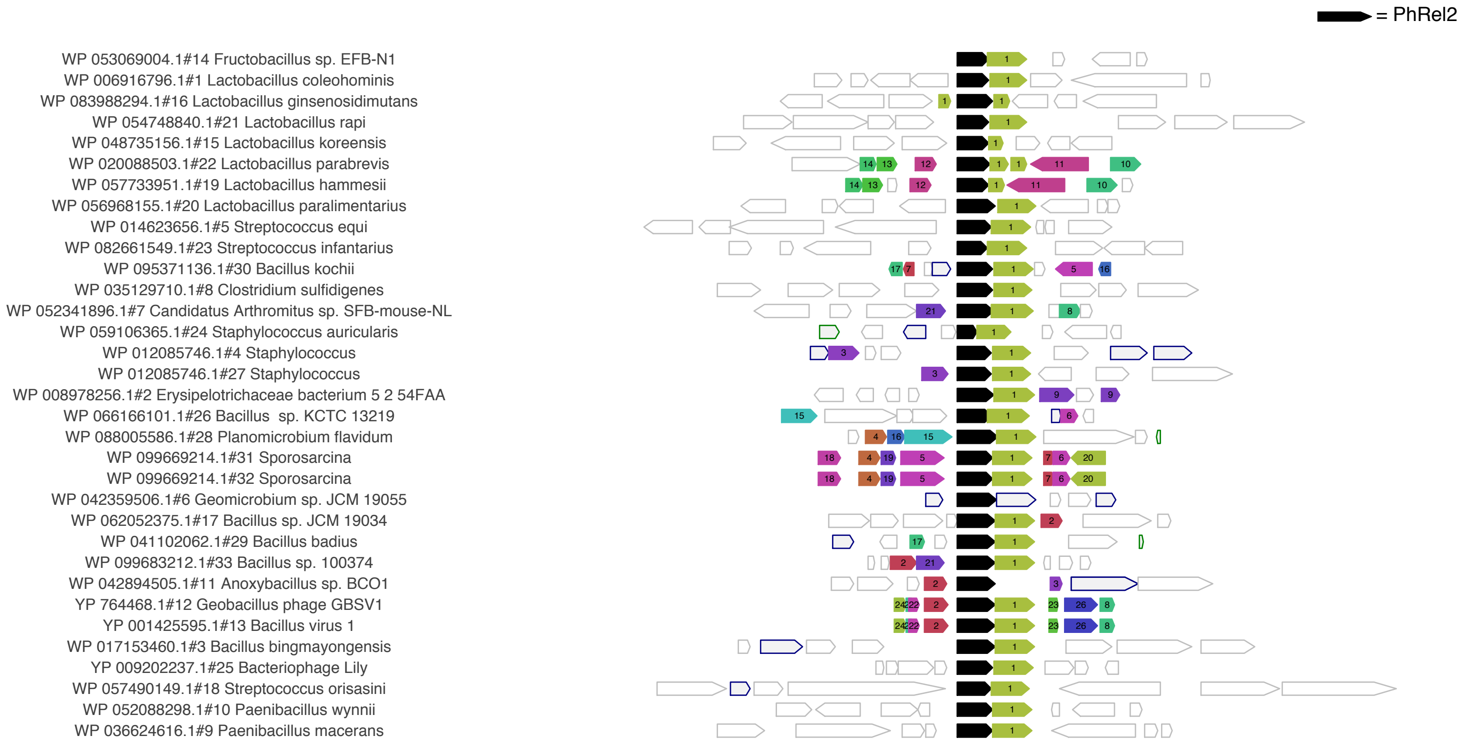

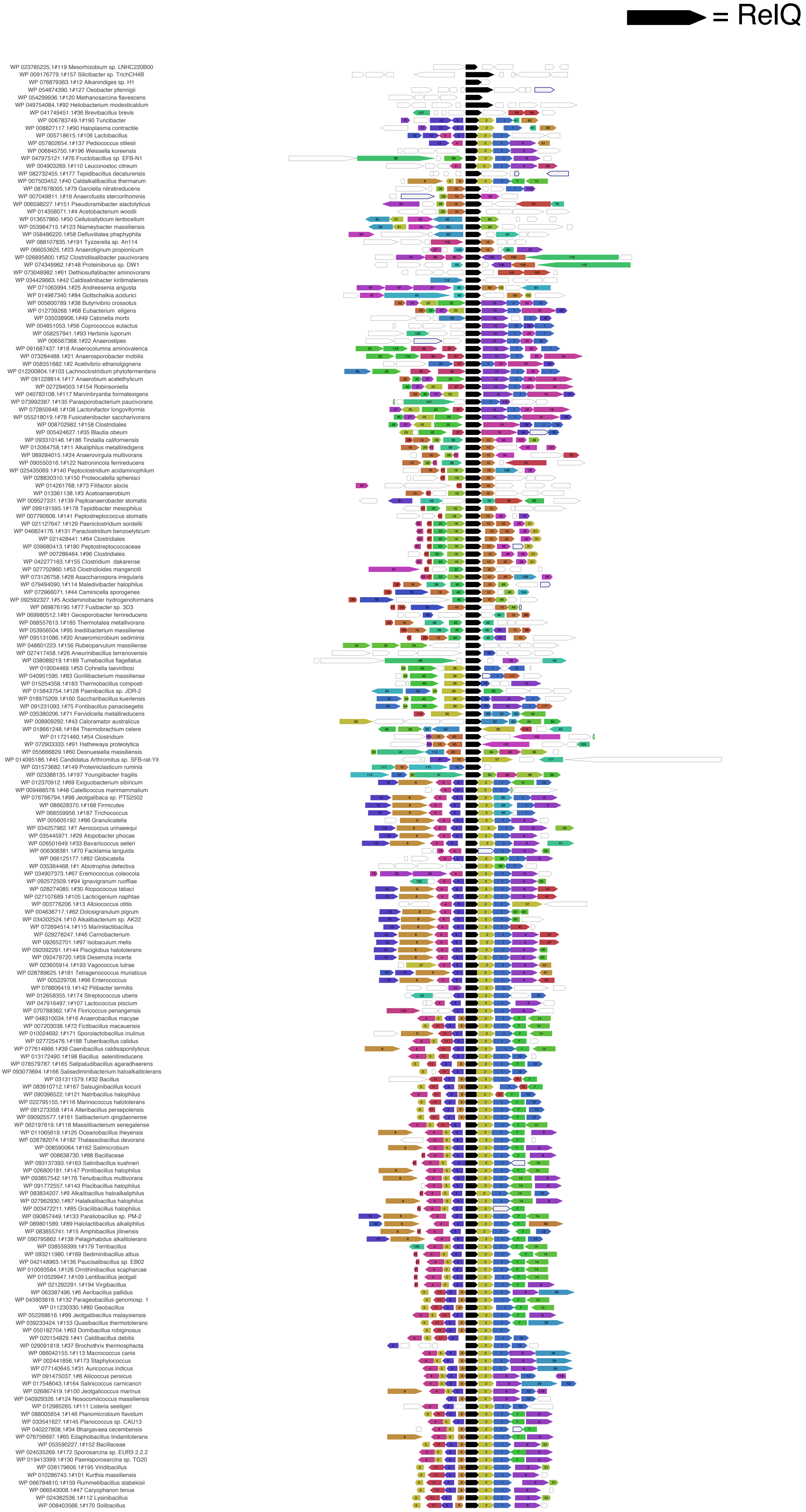

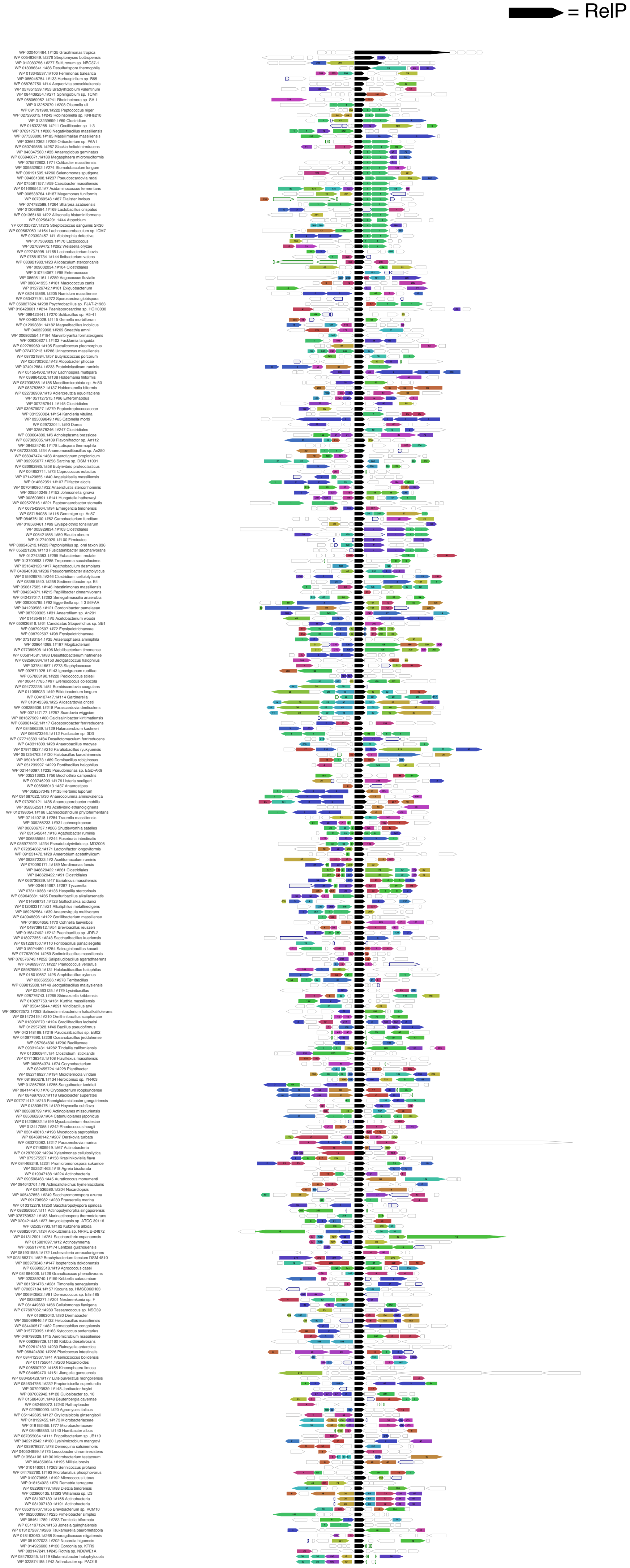

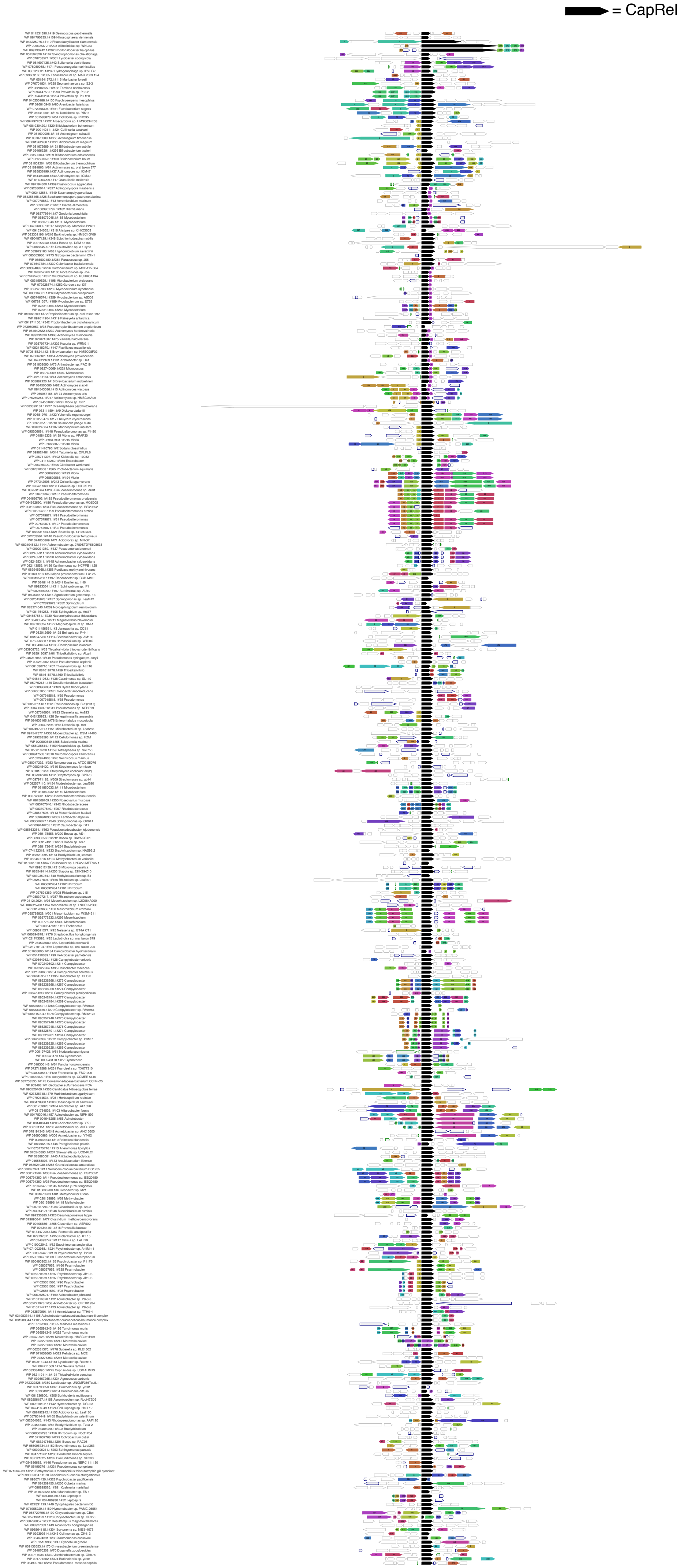

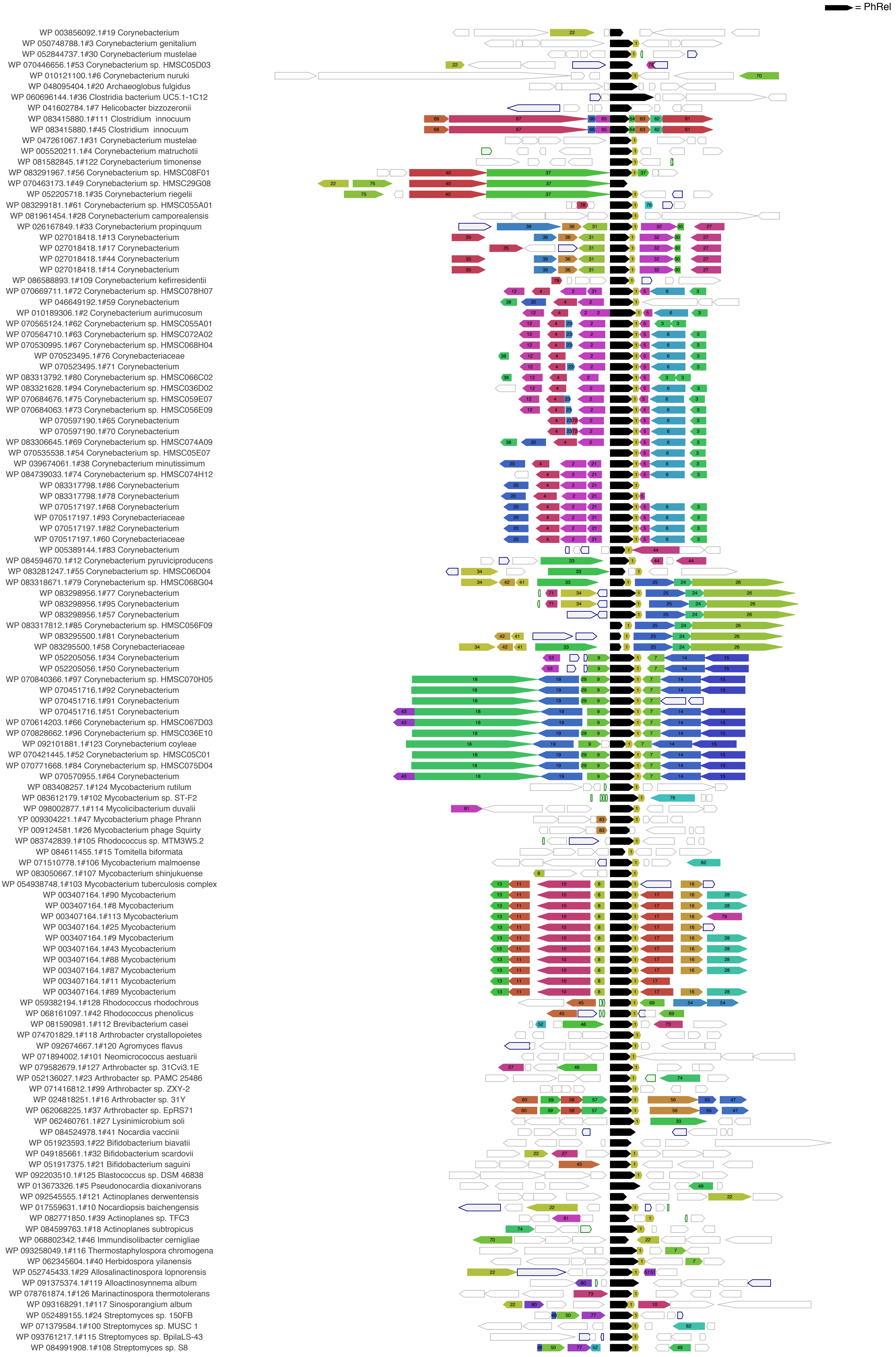
